# Amyloid-motif-dependent tau self-assembly is modulated by isoform sequence context

**DOI:** 10.1101/2023.12.13.571598

**Authors:** Sofia Bali, Ruhar Singh, Pawel M. Wydorski, Nico E. Van Nuland, Aleksandra Wosztyl, Valerie A. Perez, Dailu Chen, Jie Chen, Josep Rizo, Lukasz A. Joachimiak

**Affiliations:** Molecular Biophysics Graduate Program, University of Texas Southwestern Medical Center, Dallas, TX 75390, United States; Center for Alzheimer’s and Neurodegenerative Diseases, Peter O’Donnell Jr. Brain Institute, University of Texas Southwestern Medical Center, Dallas, TX 75390, United States; Instituto de Química Física Blas Cabrera (IQF), CSIC, E-28006 Madrid, Spain; Department of Biophysics, University of Texas Southwestern Medical Center, Dallas, TX 75390, United States Southwestern Medical Center, Dallas, TX 75390, United States; Department of Biochemistry, University of Texas Southwestern Medical Center, Dallas, TX 75390, United States; Department of Pharmacology, University of Texas Southwestern Medical Center, Dallas, TX 75390, United States

## Abstract

The microtubule-associated protein tau is implicated in neurodegenerative diseases characterized by amyloid formation. Mutations associated with frontotemporal dementia increase tau aggregation propensity and disrupt its endogenous microtubule-binding activity. However, the structural relationship between aggregation propensity and biological activity remains unclear. We employed a multi-disciplinary approach, including computational modeling, NMR, cross-linking mass spectrometry, and cell models to engineer tau sequences that modulate its structural ensemble. Our findings show that substitutions near the conserved ‘PGGG’ β-turn motif informed by tau isoform context reduce tau aggregation in vitro and cells and can even counteract aggregation induced by turn destabilizing disease-associated proline-to-serine mutations. Engineered tau sequences maintain microtubule binding and explain why 3R isoforms exhibit reduced pathogenesis compared to 4R. We propose a simple mechanism to reduce the formation of pathogenic species while preserving biological function, thus offering insights for therapeutic strategies aimed at reducing tau protein misfolding in neurodegenerative diseases.

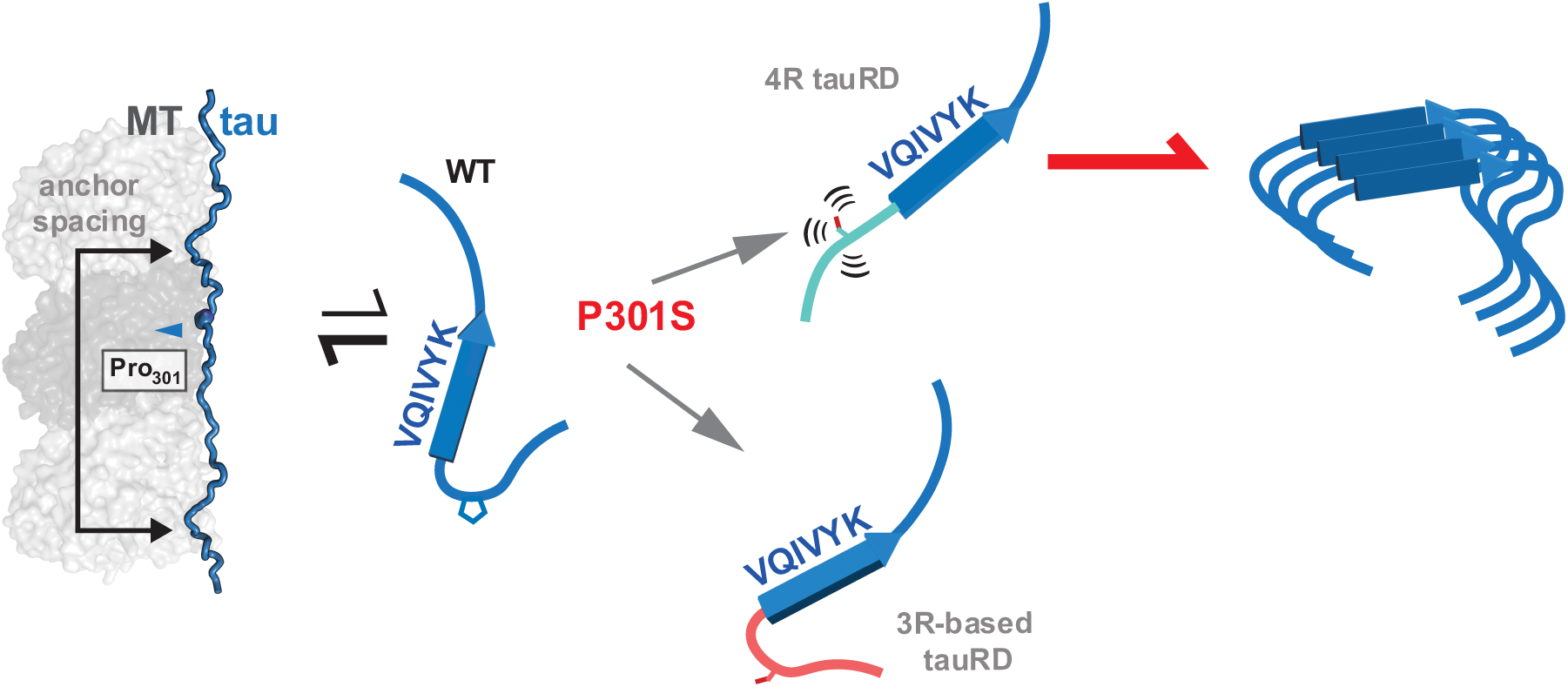

## INTRODUCTION

The deposition of the microtubule-associated protein tau is linked to a group of neurodegenerative diseases collectively known as tauopathies^1–3^. Recent cryo-EM tau fibril structures derived from patient material have uncovered how tau adopts distinct conformations (i.e., structural polymorphs) across different tauopathies^4–6^, each able to propagate in a prion-like manner^7,8^. Despite these insights into the fate of tau protein, the mechanism by which it misfolds into different conformations in each disease is poorly understood. Prior studies have shown that the normally aggregation-resistant tau monomer can convert into an aggregation-prone conformation, termed a seed, which can then recruit inert tau monomers to form amyloid fibrils^9^. These seeds have been detected in young tauopathy mouse models before the detection of insoluble species^10^ and can be isolated from Alzheimer’s disease patient material^11^. Structural and biochemical studies comparing inert tau monomers and monomeric seeds have identified rearrangements of long-range charge complementary and short-range hydrophobic interactions mediated by amyloidogenic motifs^12^.

Mutations in the *MAPT* gene that encodes tau are linked to frontotemporal dementia (FTD)^13,14^, and these mutations primarily localize to the repeat domain of tau (tauRD)^15,16^. This domain comprises the region required for binding to microtubules (MT)^17^. Alternative splicing of the *MAPT* gene results in two isoforms of the tau repeat domain: 4R tauRD (243-380) and 3R tauRD (243-380, ΔR2:275-305)^18^. Notably, 4R tau isoforms aggregate with a higher propensity than 3R tau^19^. Tau amyloid deposits in tauopathies are mostly composed of either a mixture of 4R and 3R isoforms or 4R-only isoforms, with Pick’s disease being the only tauopathy exclusively linked to the 3R tau isoform^1,20^. Experimental evidence from mouse models provides compelling insights into this isoform dynamics: reducing 4R tau isoforms reduces tau pathology and toxicity, whereas reducing 3R isoforms increases tau pathology^19^. This differential behavior of 3R and 4R tau isoforms presents avenues for therapeutic intervention, with current strategies exploring the use of antisense oligonucleotides^20,21^. A mechanistic understanding of how these isoforms alter tau’s intrinsic aggregation properties will be crucial in developing novel regulatory strategies.

A serine or leucine mutation at proline position 301, which localizes to the ^301^PGGG^304^ motif at the 3R/4R splice junction site, causes FTD^16^ and is commonly used in animal, cellular, and in vitro models to study tau aggregation^22–26^. This mutation is also upstream of the ^306^VQIVYK^311^ amyloid motif, which forms essential interactions in all tau fibril structures^27,28^, while mutations in this motif abrogate tau aggregation^11^. Moreover, time-resolved cryo-EM analysis of disease-specific tau folds has uncovered early fibrillar intermediates stabilized by VQIVYK-VQVIYK interactions^29^. Our previous studies using minimal fragments have established that sequences from both 4R and 3R tau isoforms upstream of the ^306^VQIVYK^311^ motif are sufficient to regulate amyloid motif aggregation. We observed that these sequences, which are upstream of the ‘PGGG,’ can transiently engage with ^306^VQIVYK^311^, and their interaction depends on the ‘PGGG’ sequence adopting a β-turn^24,30,31^. However, the P301S mutation from 4R tau can disrupt this aggregation inhibition, while the equivalent P270S mutation from 3R tau does not. The flexible nature of the ‘PGGG’ sequence enables the upstream sequence to adopt a range of conformations in tauopathy fibrils from disordered to folding onto the amyloid motif into different beta-arch geometries. Post-translational modifications in proximity to the amyloid motif have been shown to stabilize amyloid motif-mediated self-association into fibrils^32^. These findings underscore the complexity of the structural ensembles sampled by 4R vs. 3R tau isoforms that may balance protective anti-aggregation and pro-aggregation interactions involving amyloid motifs^32–35^.

Thus, a deeper investigation into how to regulate ^306^VQIVYK^311^–based tau aggregation will yield new insights into tuning tau ensembles to not only inhibit assembly but also to control tau folding into defined amyloid conformations. This study uses a multidisciplinary approach to understand how local sequence changes modulate the ‘PGGG’ structural motif to limit amyloid motif-dependent assembly. By leveraging the natural sequence composition of the 3R isoform, we evaluate the structural and chemical impacts that reduce aggregation while maintaining microtubule-stabilizing function. Our research presents a model for inhibiting 4R tau aggregation, potentially opening new avenues to develop targeted therapeutic strategies for tauopathies.

## RESULTS

### Differential aggregation propensity induced by sequence preceding the amyloid motif ^306^VQIVYK^311^ in 3R vs. 4R tau

The 4R and 3R tau isoforms differ in sequence upstream of the ^306^VQIVYK^311^ motif at only five out of 11 amino acids (Fig. 1a). We compared ThT-based aggregation propensities of wild-type and FTD-linked proline to serine mutants in 4R and 3R peptides that span the inter-repeat region to repeat 3 (Fig. 1b-c and Supplementary Fig. 1a-d). We determined that the P301S R2R3_295-311_ peptide assembled into amyloid fibrils rapidly starting at 100 μM; peptide WT R2R3_295-311_ only aggregated at the highest concentration, 800 μM, while peptides WT R1R3_295-311ΔR2_ and P270S R1R3_263-311ΔR2_ did not aggregate at any concentration within the time frame of the experiment (Fig. 1c and Supplementary Fig. 1a-d). We evaluated the aggregation behavior of WT and P301S 4R tauRD (residues 244-380), as well as WT and P270S 3R tauRD (residues 244-380Δ275-305) (Supplementary Fig. 1e). These longer tau fragments replicated the differential proline to serine aggregation phenotypes observed in the P270S R1R3_263-311ΔR2_ and P301S R2R3_295-311_ peptides.

**Figure 1.**
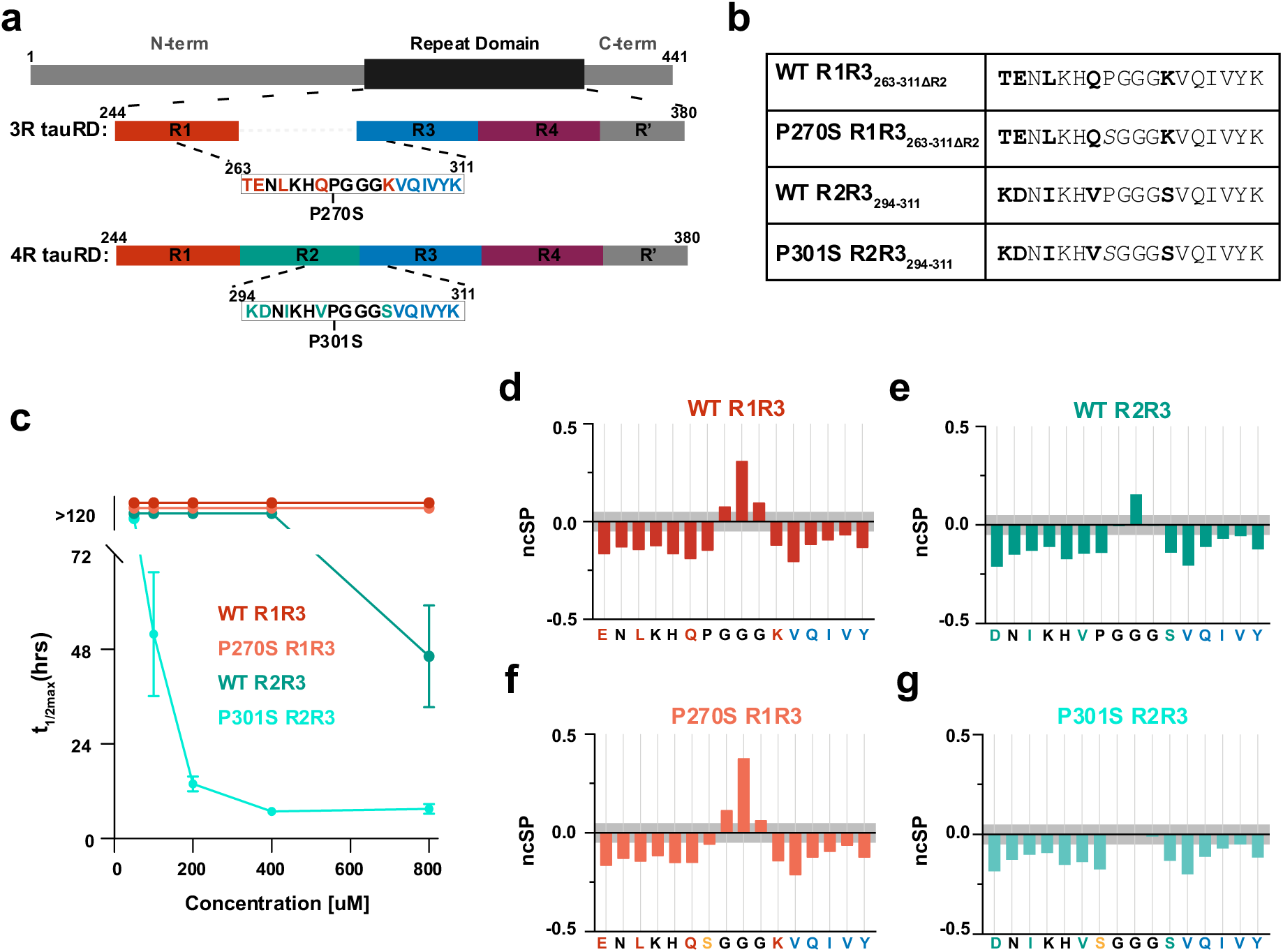
Proline to serine mutations in a 4R-like fragment leads to a more unstructured turn. **a.** Cartoon schematic of 3R and 4R-like tau fragments. Repeat domains are colored red, green, blue, and purple of repeats 1, 2, 3, and 4. Inset shows WT minimal fragments that span repeats 1 and 3 and repeats 2 and 3. The site of the proline to serine mutation is indicated below the sequence. **b**. Table listing the four tested peptide sequences. **c**. ThT fluorescence aggregation assay of WT R1R3_263-311ΔR2_ (red), P270S R1R3_263-311ΔR2_ (orange), WT R2R3_295-311_ (green), and P301S R2R3_295-311_ (cyan) peptides carried out at 5 concentrations: 50, 100, 200, 400, and 800 µM. Data is shown as the t_1/2max_ fit to each aggregation curve. The data is plotted as an average with a standard deviation. **d-g**. neighbor corrected structure propensity (ncSP) calculated from peptide 25°C Hα chemical shift assignments using ncSPC calculator for WT R1R3_263-311ΔR2_ (d, red), P270S R1R3_263-311ΔR2_ (e, green), WT R2R3_295-311_ (f, orange), and P301S R2R3_295-311_ (f, cyan). Negative/positive values correspond to β-sheet/α-helical propensity. Gray bars highlight the deviations expected for random coil structures.

We acquired TOCSY and NOESY spectra for the four peptides at 25°C and 10°C to investigate the local structural differences sampled by these four sequences (Fig 1b.). We performed proton chemical shift assignments for each sequence at 10C° and transferred assignments to 25°C (Supplementary Fig. 2a-d). For the aggregation-prone peptide, P301S R2R3_295-311_, we confirmed that signal intensities did not significantly change throughout the experiment at 25°C (Supplementary Fig. 1f). We observed small deviations when comparing the chemical shifts of WT R1R3_263-311ΔR2_ to P270S R1R3_263-311ΔR2_ and WT R2R3_295-311_ to P301S R2R3_295-311_ except for the expected shifts at the position preceding the proline to serine mutation (Q269 and V300 respectively, Supplementary Fig. 2a,b and Supplementary Table 1). We estimated the extent of b-sheet or α-helical content in our fragments based on Cα proton chemical shifts (Hα) using the neighbor-corrected secondary structure propensity calculator (ncSPC) with a neighbor correction window of 3^36,37^.

All four peptides exhibited a similar secondary structure profile at 25°C and 10°C with more negative ncSP values at 25°C (Fig. 1d-g; Supplementary Fig. 1g-j). The profiles showed negative scores consistent with an enhanced population of β-sheet structure compared to random coil for the left and right side of the ‘PGGG’/’SGGG’ motif (Fig. 1d-g). The central ‘PGGG’ motif (270-273) in WT R1R3_263-311ΔR2_ had a higher turn propensity than the motif (301-304) in WT R2R3_295-311_. In the peptides containing the mutant ‘SGGG’ motif, P270S R1R3_263-311ΔR2_ had a similar turn propensity as its proline-containing counterpart. In contrast, the P301S R2R3_295-311_ peptide showed no significant deviations from the random coil signal (Fig. 1d-f). Therefore, we observed subtle differences localized to the ‘PGGG’/’SGGG’ motif specific to the P301S R2R3_295-311_ peptide.

A simple hypothesis is that the R1R3 sequence is less compatible with amyloid fibrils than the R2R3 sequence. To test this, we assessed the role of these sequences in the end-stage tau fibrils derived from tauopathy patients (Supplementary Fig. 1k). We observe that these sequences adopt a large variety of conformations within the fibril form: dynamic and unresolved in the fibril structure (AD/CTE), forming a beta arch that interacts on either the polar (“-Q-V-K” GGT/CBD/AGD,) or nonpolar (“V-I-Y” GPT/PSP,P301Sm) face of the amyloid motif, and only in the 3R derived fibril is this segment extended (PiD; Supplementary Fig. 1k). Comparing this segment to tau P301S fibrils extracted from a tauopathy mouse model, we observe that the mutant segment adopts yet a different beta-arch conformation that folds onto the “V-I-Y” face (Supplementary Fig. 1k). Overlaying these regions, we observe that the ‘PGGG/SGGG’ forms a pivot point where the flexibility of this motif enables the access of interactions between both faces of the ‘VQIVYK’ motif with different residues in the 294-305 element, with both the Proline and Serine at position 301. To evaluate the individual contributions of each residue, we compared the solvent-accessible surface area (SASA) to the estimated contribution of each residue to the fibril structure through an alanine^27^ (Supplementary Fig. 1m-n). As previously described^27^, we observe that the VQIVYK motif is buried and is important to the stability of the assembly (Supplementary Fig. 1m-n). In the AD and CTE tau folds, the sequence starts at the amyloid motif, and the preceding sequences are not ordered (Supplementary Fig. 1k). In the other structures, the upstream residues from R2 (D295, I297, V300, and S305) and R1 in the PiD structure (E264, L266, Q269, and K274) are less buried than the VQIVYK amyloid motif (Supplementary Fig. 1m). Still, their contribution to stability is variable (Supplementary Fig. 1n). Of all the residues upstream of VQIVYK, I297 in R2 or L266 in R1 appear to have the largest and most consistent contribution to the local stability of tau fibrils with the leucine and isoleucine residues largely isosteric (Supplementary Fig. 1k, m-n) except for the P301S tau structure where the Cγ of I297 forms a clash. Similarly, D295 in R2 and E265 in R1 are either buried and form salt bridges or face the solvent (Supplementary Fig. 1k, m-n). Finally, the residues that flank the PGGG motif are buried and may be more sensitive to mutations of valine to glutamine at 300 and serine to lysine at 305 from R2 to R1 (Supplementary Fig. 1k, m-n). From these analyses, we predict that for the CBD and AGD tauopathy conformations, the 3R sequences would be incompatible with the beta-arch conformations. At the same time, they should be compatible with AD, CTE, PiD, and likely also PSP, GGT, and GPT. This partitioning of sequence compatibility follows disease classification based on tau isoform pathology that is mixed 3R/4R for AD and CTE, 3R only for PiD, and 4R only for the others. Thus, the sequence upstream of VQIVYK in 3R tau (i.e., R1R3) may not be able to adopt specific beta-arch conformations, yet it is largely compatible with the available tau fibril structures. Therefore, understanding how the changes in local dynamics of this region in the monomeric state may uncover the role of the pre-fibril state of this region in defining how to regulate tau folding into amyloids.

### Molecular Dynamics ensembles differentiate the ‘SGGG’ structure of the aggregation-prone sequence

We employed Molecular Dynamics (MD) simulations of these peptide model systems to assess differences in intramolecular interactions and accessible states. The MD simulations for the four peptides (Fig 1b.) were performed using a two-stage production scheme followed by analysis with a Markov State Model (Supplementary Note 1 and Supplementary Fig. 3). Simulations were generated using canonical sampling schemes with the Amber-991lsb ff which produces ensembles suitable for disordered sequences, but with a bias towards increased β-sheet propensity^38,39^ (Fig. 2a). From the resultant 15µs of final production simulations we calculated the distance between the backbones of the C-terminal amyloid motif ‘VQIVYK’ and the N-terminal residues ‘TENLKH’ or ‘KDNIKH’ and observed wider distributions for the proline-to-serine peptides with higher sampling of states with backbone distances between 0.5-1.0 nm for P301S R2R3_295-311_ (Fig. 2b). Furthermore, we assessed the relative flexibility for each residue across the simulation using the root mean squared fluctuation (RMSF). We observed that the four peptides had relatively similar fluctuation, with only a significant increase in flexibility for residues in the ‘SGGG’ motif of P301S R2R3_295-311_.

**Figure 2.**
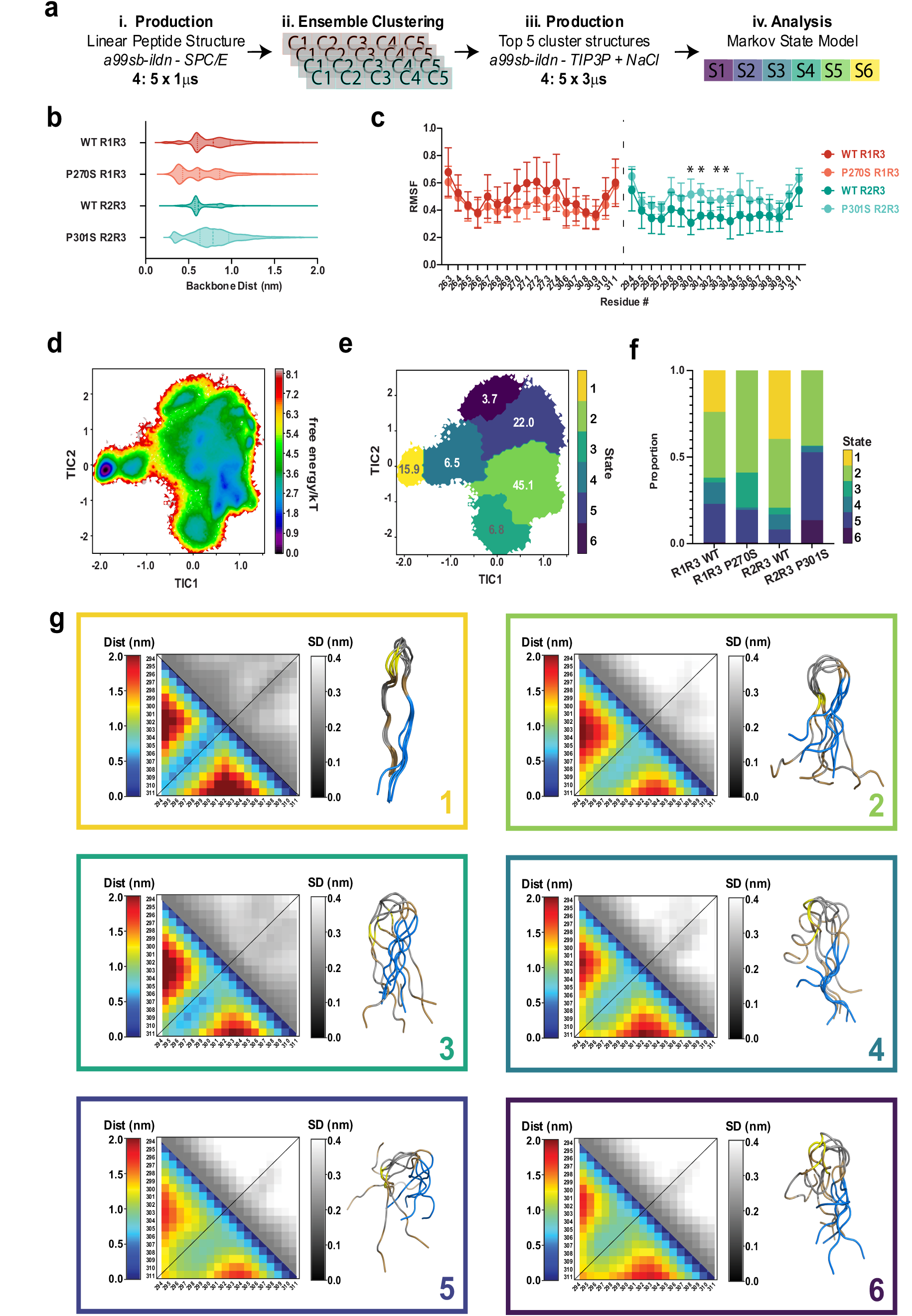
Building a structural ensemble of tau fragments reveals a distinct behavior of the aggregation-prone 4R-derived peptide. **a.** Two stage MD simulation workflow for: WT R1R3_263-311ΔR2,_ P270S R1R3_263-311ΔR2,_ WT R2R3_295-311,_ and P301S R2R3_295-311_ **b.** backbone distance between the center of mass of the N-term residues 1-6 (TENLKH/KDNIKH) and C-term amyloid motif residues (VQIVYK) second-stage MD ensembles. The data is shown as violin plots with median and 25/75% quartiles labeled in dash lines. **c.** Per-residue RMSF of 3R-derived (left) and 4R-derived (right) peptides from second-stage MD ensembles. The data is shown as average +/− SD across the 5 repeat simulations. **d.** Free Energy landscape mapped onto the first two TICA components calculated from MD ensemble MSM for the combined simulations of WT R1R3_263-311ΔR2_, WT R2R3_295-311_, P270S R1R3_263-311ΔR2_, and P301S R2R3_295-311_ for a total of 60µs. Data is colored based on normalized free energy from black to red. **e.** MSM metastable state map, states 1-6 mapped from yellow to purple on the free energy landscape. **f.** Bar plot illustrating the proportion of sampling of each state per peptide. The states sampled for each peptide are colored in c. **g**. State-specific Average(bottom) and Standard deviation (top) maps of pairwise distance calculated for all timepoints in each state in boxes colored in c. Average (bottom) colored from blue to red, 0 nm to 2.0 nm. Standard deviation (top) colored from black to white, 0 nm to 0.4 nm. The first 5 structures per state are shown in cartoon representation with ‘VQIVYK’ colored in blue, unique residues colored in brows, Proline/Serine position colored in yellow, and similar residues colored in gray.

We built and validated a Markov State Model (MSM) with pyEMMA^40^ (see Supplementary Note 1 and Supplementary Fig. 3). From the MSM, we visualized a free energy surface and mapped the 6 metastable states of the landscape (Fig. 2d-e). For each state, we calculated the relative frequency for each peptide and generated heatmaps for the mean distance (rainbow) and deviation (grayscale) using the 4R isoform numbering (Fig. 2f-g, states 1-6). We observed that the lowest energy state (Fig. 2e-g, state 1, yellow) was populated only by the ‘PGGG’ turn-containing peptides (WT R1R3_295-311ΔR2_ and WT R2R3_295-311_) and is composed of defined β-hairpin structures with contacts above the diagonal between residues 295-299/305-309 (Fig. 2g, state 1). The next lowest energy state (Fig. 2e-g, state 2, green) was sampled by all four peptides and represents a collection of local minima with structures forming a less defined turn and a shifted register with contacts below the diagonal (Fig. 2g, state 2). P270S R1R3_295-311ΔR2_ primarily populated state 3 and had contacts between 297-299/305-309 (Fig. 2e-g, state 3, turquoise). State 4 (Fig. 2e-g, state 4, teal) was primarily populated by WT R1R3_295-311ΔR2_ and WT R2R3_295-311_ and represents an interchange from the local minima state 1 and the rapidly exchanging state 2 with interactions below the diagonal (Fig. 2g, state 4). Higher energy state 5 (Fig. 2e-g, state 5, indigo) is also accessible to all four peptides but is preferentially sampled by the P301S R2R3_295-311_ sequence representing a more expanded structure with medium range interactions below the diagonal (Fig. 2g, state 5). By contrast, state 6 was uniquely populated by the P301S R2R3_295-311_ defined by transient interactions above the diagonal, leaving reduced contacts to ^306^VQIVYK^311^ (Fig. 2e-g, state 6, purple).

These observations reveal a unique sampling of states across the four peptides. Sequences with P301 at the ‘**PG**GG’ motif promote sampling of a β-hairpin-like structure with transient interactions across the peptide sequence. The proline to serine mutation disrupts the turn localization. However, the sequence properties of ‘TENLKHQ-K’ in P270S R1R3_295-311ΔR2_ increase the sampling of states with a left-shifted turn ‘SG**GG**’ that still contains medium-distance contacts to VQIVYK. In the P301S R2R3_295-311_ peptide, the ‘DNIKHV-S’ sequence shifts sampling to states with a turn centered before the ‘SGGG’ motif, and the induction of a hydrophobic residue at position 300 increases the sampling of interactions to residues 305-307 resulting in reduced interactions to the latter part of the VQ**IVYK** motif.

### Ensemble-based design of 4R tauRD mutants reduces tau aggregation propensity

To design aggregation-resistant 4R tauRD, we based our strategy on the sequence differences from the 3R isoform and incorporated them in a 4R context. The first construct uses the sequence from R1 with V300Q and S305K mutations (herein, tauRD_QK). The second mutant incorporates the full R1 element with four mutations: D295E, I297L, V300Q, and S305K (herein, tauRD_EL_QK). In addition to testing the constructs with the wild-type proline (i.e., ‘PGGG’ motif), we tested our designs in the context of the P301S mutation (i.e., ‘SGGG’ motif) (Fig. 3a). We evaluated the aggregation propensity of the constructs in a ThT fluorescence aggregation assay induced with heparin. We fit the resultant curves to estimate a t_1/2max_ (Supplementary Fig. 4a-f). The presence of fibrils at the endpoint of the aggregation reactions for the 10 µM and 40 µM reactions was confirmed by TEM (Supplementary Fig. 4g, h). We observed that the tauRD_QK mutant aggregated with similar rates to WT tauRD (Fig. 3b and Supplementary Fig. 4a, b). By contrast, the tauRD_EL_QK showed values that were 2- and 3-fold slower than WT tauRD (Fig. 3b and Supplementary Fig. 4c). The P301S tauRD_QK aggregated between 2- and 4-fold more slowly compared to P301S tauRD while the P301S tauRD_EL_QK further reduced aggregation leading to t_1/2max_ values between 3- and 5-fold slower than P301S tauRD (Fig. 3c and Supplementary Fig. 4d-f).

**Figure 3.**
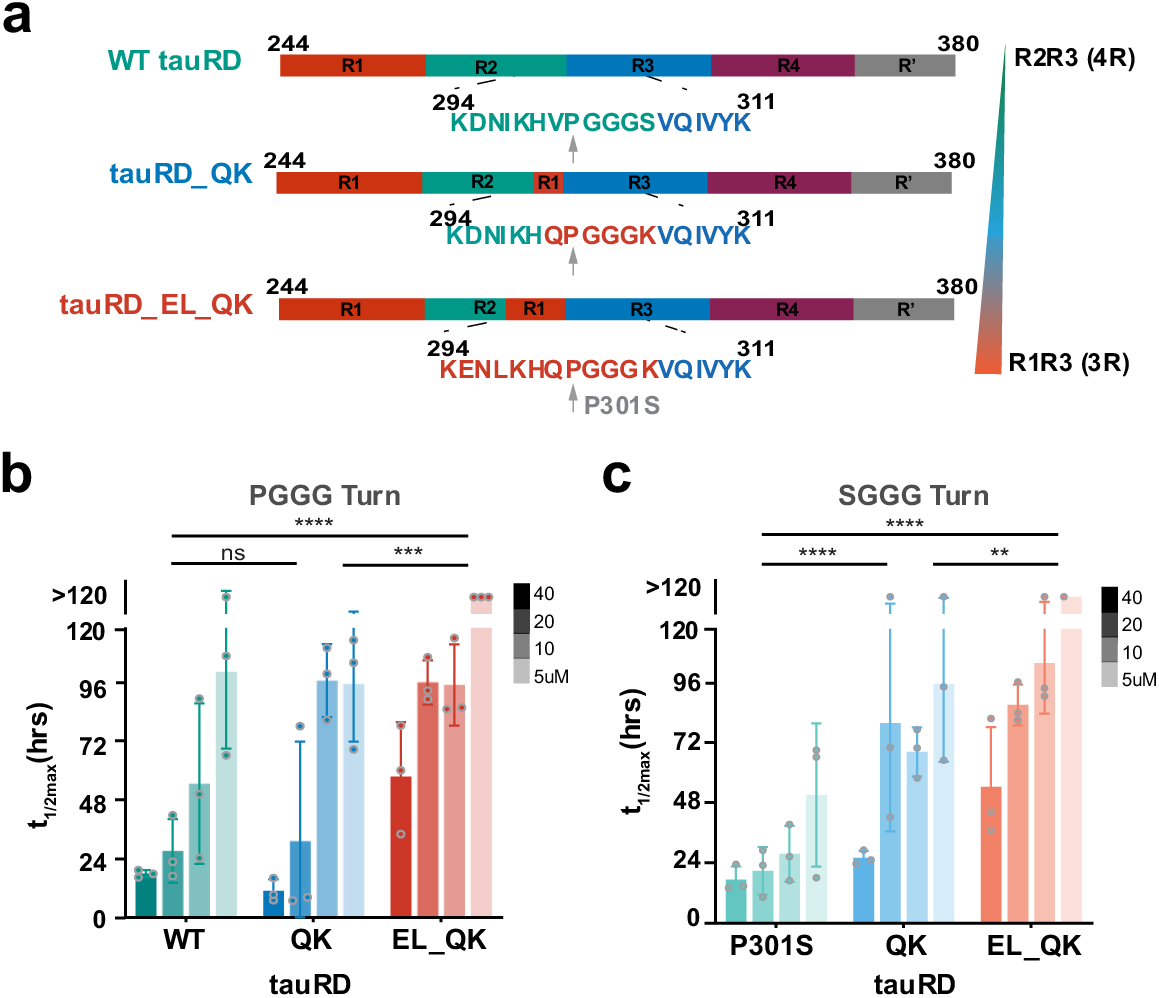
Designed tau sequences reduce aggregation of tau in vitro. **a.** Cartoon schematic of 4R tauRD and designed tauRD_QK and tauRD_EL_QK. Repeat domains are colored as in Figure 1, and the sequence inset of designed mutations below each schematic with residues is colored based on repeat. **b-c.** ThT fluorescence aggregation assay of tauRD constructs induced with 2X heparin carried out at 4 concentrations: 5, 10, 20, 40 µM (light to dark) for ‘PGGG’ containing constructs WT tauRD (green), tauRD_QK (blue), and tauRD_EL_QK (red) in b, and ‘SGGG’ containing constructs P301S tauRD (cyan), P301S tauRD_QK (sky blue), and P301S tauRD_EL_QK (orange) in c. Data is shown as the t_1/2max_ fit to each aggregation curve. The data is plotted as an average with a standard deviation, significance calculated from column effects using 2way ANOVA (*=p < 0.05, **=p < 0.01, ***=p < 0.001, ****=p < 0.0001).

Additionally, we evaluated our selected mutations in the context of 3R tauRD (244-380Δ275-305) using the reverse mutations that include E264D, L266I, Q269V, and K274S (herein, 3R tauRD_DI_VS) in constructs encompassing the full sequence observed in the ex-vivo Pick’s disease fibril structure^41^. ThT fluorescence intensity endpoints were used to evaluate the heparin-induced aggregation propensity of the constructs, and the fibrils were visualized by TEM (Supplementary Fig. 4i-k). 3R WT tauRD and 3R tauRD_DI_VS showed baseline ThT signal and no fibrils by TEM. By contrast, the 3R P270S tauRD had a reduced ThT signal with minimal fibrils observed by TEM relative to the 4R P301S tauRD construct (Supplementary Fig. 4j). However, 3R P270S tauRD_DI_VS increases fibril formation and produces ThT signal at about 70% of the 4R tauRD P301S (Supplementary Fig. 4j). Therefore, we found we could increase the aggregation sensitivity of the 3R tauRD by incorporating the sequence from repeat 2 but in the context of a destabilized ‘SGGG’ turn. Taken together, we observe that introducing the 3R-like mutations into 4R tauRD near the VQVIYK amyloid motif reduces the aggregation propensity of tau.

### Balance of intermediate and long-range contacts correlates with inhibition of tau aggregation

To understand how the highly dynamic intramolecular interactions in tauRD change in response to the designed mutations, we turned to XL-MS. This approach enables the capture of interactions in complexes to define the architectures of assemblies^42–44^. More recently, our lab has established ways to probe changes in the conformation of IDPs using XL-MS^12,30,45^. Previously, we have used this approach to probe the effect of P301S on tauRD dynamics, uncovering how this mutant is more sensitive to unfolding and, thus, more competent to self-associate^30^. Here, we extend this approach to understand how the designed mutations change the distribution of local contacts in proximity to the mutation and longer-range contacts to relate these changes to the reduction of tau aggregation. The 6 purified tauRD constructs (Fig. 4a) were reacted with the lysine-reactive DSS cross-linker (DSS, spacer arm 11.4A). The samples were analyzed by mass spectrometry, and the data was processed using the xQuest pipeline^46^. Each sample was processed as 5 independent replicates, and only consensus contacts were considered for further analysis (Fig. 4a).

**Figure 4.**
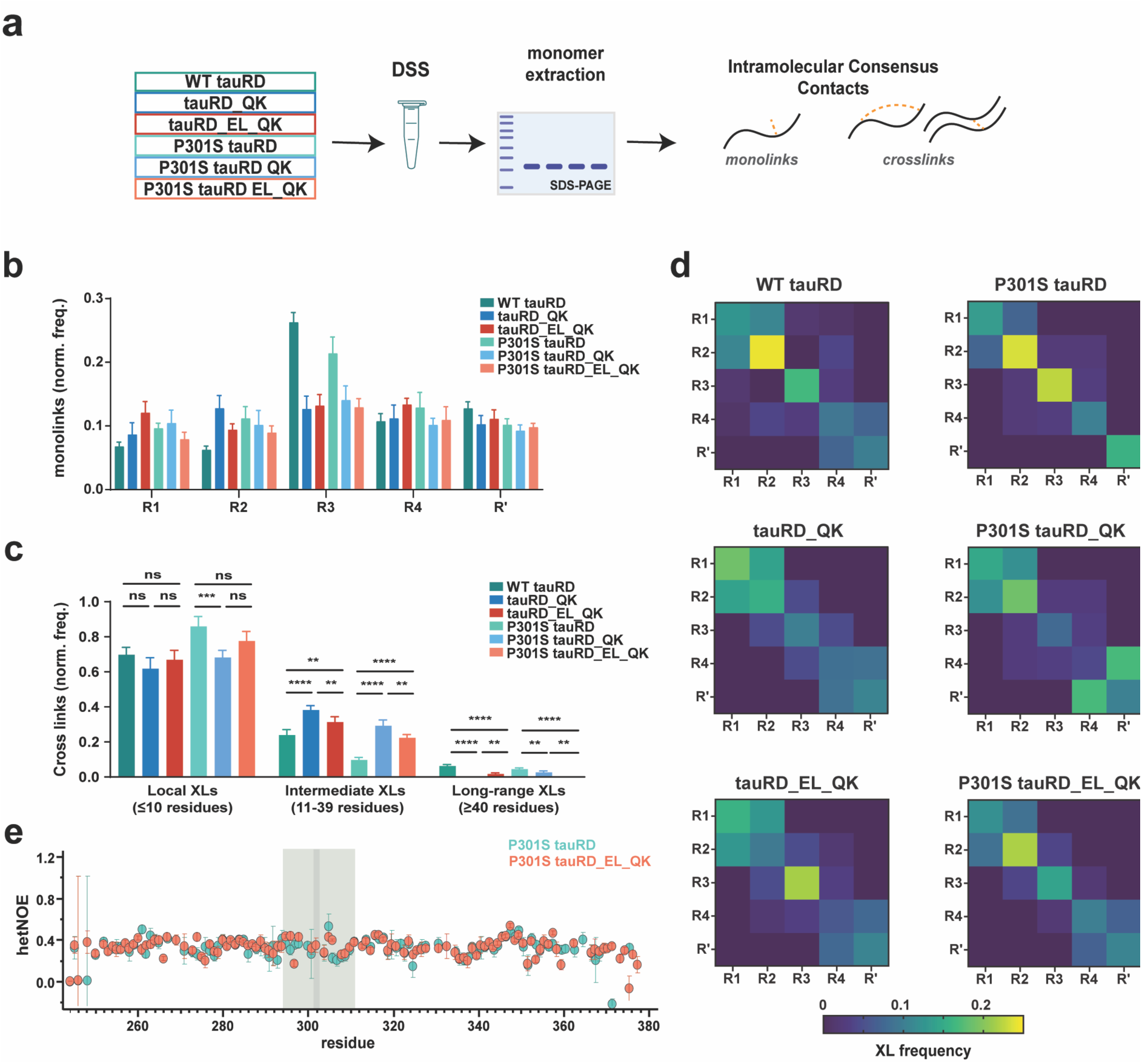
3R-based mutations increase intermediate-range interactions. **A.** Schematic of sample preparation for XL-MS workflow. Tau constructs: WT tauRD (green), tauRD_QK (blue), and tauRD_EL_QK (red), P301S tauRD, P301S tauRD (cyan), P301S tauRD_QK (sky blue), and P301S tauRD_EL_QK (orange) were cross-linked with disuccinimidyl suberate (DSS) then ran on an SDS-PAGE gel for monomer extraction, digestion, and purification to be then processed through our XLMS workflow for identification of consensus mono-links, and loop-link/cross-links. **b.** Quantification of identified mono-links grouped by repeat: R1 (244-274), R2 (275-305), R3 (306-336), R4 (337-368), and R’ (369-380); for tauRD constructs (colored as in a). Data is shown as the sum of the normalized mean and standard deviation across 5 replicated of the nseen for each construct. **c**. Quantification of local (< 10 residues), intermediate (11-39 residues), and long-range loop-links/cross-links (> 40 residues) by sample (colored as in a). Data is plotted as an average with a standard deviation, significance calculated one way ANOVA (*=p < 0.05, **=p < 0.01, ***=p < 0.001, ****=p < 0.0001). **d**. Heatmap of distribution of loop-links/cross-links for each construct grouped by repeat as in c. Data is shown as the average normalized frequency of nseen for each construct colored from navy to yellow. **e.** ^15^N-^1^H heteronuclear NOE experiments for P301S tauRD (cyan) and P301S tauRD_EL_QK (orange). The ratio is calculated by comparing the integrated peak volume between the saturation and reference spectra (see Supplementary Fig. 6f). Peak errors were determined from the standard deviation of the intensity distribution in signal-less spectral regions. 295-311 region marked by the green bar, G302 residue marked with a gray bar.

To look at the relative accessibility of regions in tauRD, we determined the normalized frequency of mono-link modifications, defined as DSS modifications on a single lysine, mapped onto each repeat. WT and P301S tauRD show significantly higher frequencies of monolinks in repeat 3 despite a similar frequency of modification across the rest of the sequence (Fig. 4b and Supplementary Fig. 5a). Next, we mapped the loop-links/cross-links (XL) frequency in these samples, defined as intramolecular covalent contacts between pairs of lysines located on one or two peptides, showing unique cross-link pair colored by the normalized frequency (Supplementary Fig. 5b). We grouped and clustered these XL by contact order: contacts within repeats (local; ≤ 10 residues), between adjacent repeats (intermediate; 11-39 residues), and further (long-range; ≥ 40 residues) (Supplementary Fig. 5c). First, a comparison of the WT and P301S tauRD analysis reaffirms our previous findings that the P301S mutation appears to destabilize intermediate and long-range contacts within the repeat domain of tau^30^, while the relative frequency of local XL increases (Fig. 4c). For our four designed mutants, we observe an increase in intermediate contacts coupled with a decrease in long-range XL and a decrease in local XL. (Fig. 4c and Supplementary Fig. 5c).

Mapping these cross-links by repeat domain (i.e., R1-R4), we observe that the shift from long-range to intermediate interactions corresponds to a shift from N-terminal interactions in Repeat 1-2 in WT and P301S tauRD to more central interactions in repeats 2-4 for the designed constructs (Fig. 4d). Finally, we relate the abundance of contacts to K311 within the VQIVY<u>K</u> amyloid motif and find that the designed constructs have more abundance of unique interactions at this position. We see interactions in our region of interest through XL at lysine positions K280, K281, and K311, which are not identified in either WT or P301S tauRD (Supplementary Fig. 5d).

To quantify residue level changes in turn dynamics proximal to the amyloid motif in WT tauRD, P301S tauRD, tauRD_EL_QK, and P301S tauRD_EL_QK, we employed ^15^N-^1^H heteronuclear NOE (hetNOE) relaxation experiments^47^. Assignments were transferred from published spectra (bmrb 19253)^48^ and supplemented with ^15^N-NOESY-HSQC and ^15^N-TOCSY-HSQC experiments to assign the mutant sites D295E, I297L, and S305K, assignment for V300Q not made (Supplementary Fig. 5e and Source Data 6). Overall, we observe high levels of dynamics across the entirety of each sequence with no significant difference in rigidity for the 295-311 region (Supplementary 6a-e). Interestingly, we observe lower signal-to-noise for P301S tauRD and P301S tauRD_EL_QK and no signals above the noise for the G302 position for P301S tauRD (Supplementary Figure 6d). Higher concentration experiments for P301S tauRD and P301S tauRD_EL_QK improve signal-to-noise for this region (Supplementary Fig. 6f), but peak intensities above the noise for position 302 in the saturation spectra of P301S tauRD were not recovered consistent with higher dynamics (Fig. 4e). The consistency in dynamics across the tauRD constructs suggests the differences that confer distinct aggregation behavior do not alter the protein dynamics globally suggesting similar ensemble diversity across the tauRD constructs.

Across the designed constructs, the most significant changes localize to intermediate-range interaction between repeats 2, 3, and 4. Overall, we find that a balance of long and intermediate contacts is correlated with aggregation inhibition, and the loss of these combined with the enhancement of localized ‘SGGG’ turn dynamics increases the unfolding and exposure of the amyloid motif.

### Aggregation-reducing tau mutations maintain microtubule-binding activity

The endogenous function of tau consists of stabilizing microtubules (MT)^17^. We performed a fluorescent MT polymerization assay coupled with molecular modeling of the MT: tau complex to determine how our engineered tau constructs alter this function. Purified tubulin dimer was incubated with each of the 6 tau constructs (i.e. Fig 3a). The rate and extent of polymerization relative to tubulin alone were determined by fitting the resultant curves (Supplementary Fig. 7a, b). For the ‘PGGG’ turn constructs, we observe that tauRD_EL_QK had increased polymerization by 1.5-fold relative to WT. In contrast, tauRD_QK polymerization amplitude was significantly reduced by 1.5-fold (Fig. 5a and Supplementary Fig. 7a). However, all three constructs had no significant difference in the t_1/2max_ time to endpoint polymerization (Fig. 5b and Supplementary Fig. 7a). Mutation of the proline to serine at position 301 has previously been shown to alter binding to MT^49,50^. Consistent with this observation, we see a nearly 10-fold reduction of polymerization for P301S tauRD compared to WT tauRD (Fig. 5a and Supplementary Fig. 7b). P301S tauRD_QK and P301S tauRD_EL_QK constructs do not significantly recover MT binding (Fig. 5a); however, the above baseline polymerization of the P301S tauRD_QK had a t_1/2max_ time approaching that of the ‘PGGG’ containing tauRDs (Fig. 5b and Supplementary Fig. 7b).

**Figure 5.**
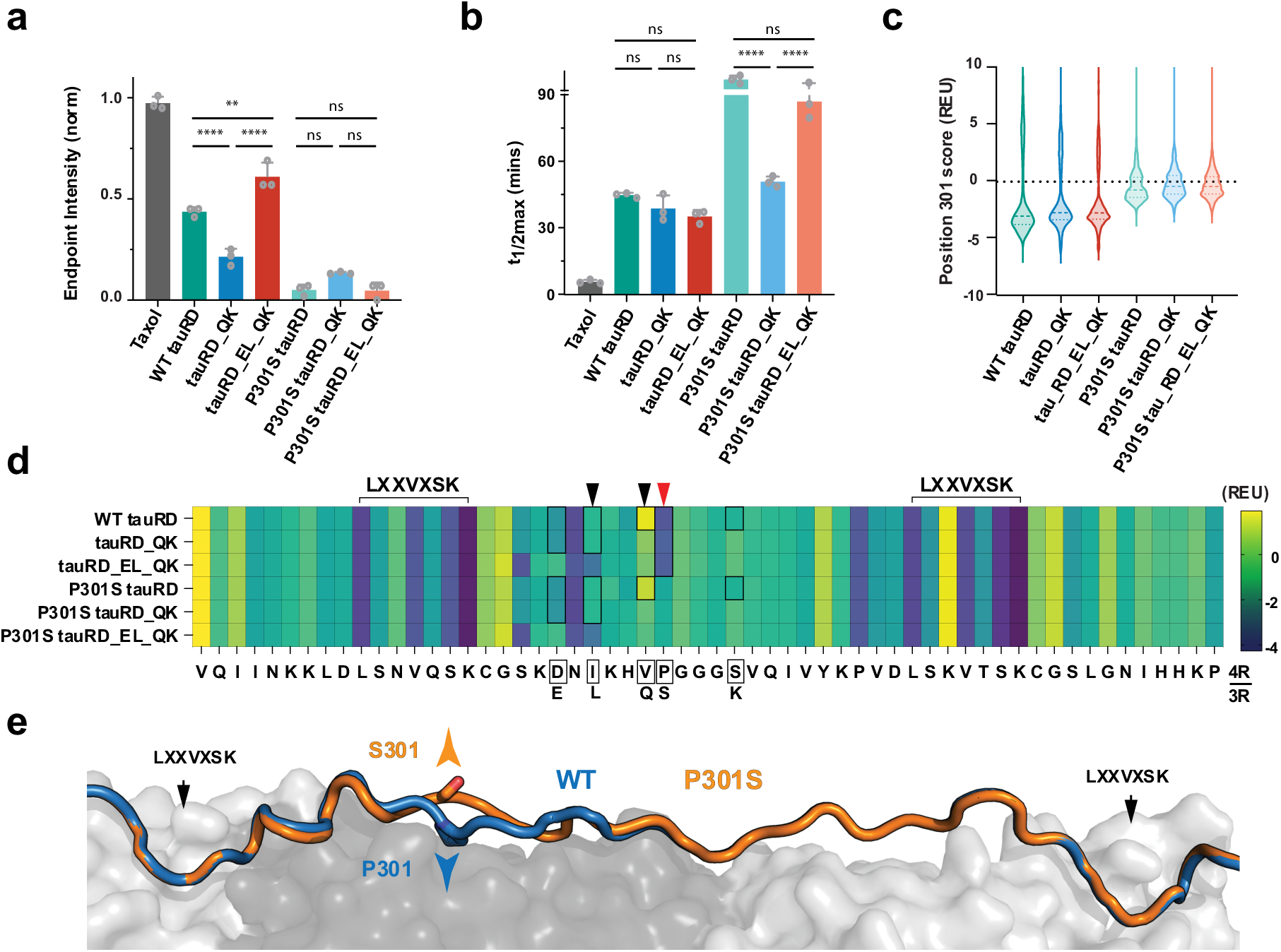
Designed tau sequences retain MT activity. **a.** Microtubule polymerization assay endpoint intensity of tauRD constructs at 10 µM for ‘PGGG’ containing constructs WT tauRD (green), tauRD_QK (blue), tauRD_EL_QK (red), and ‘SGGG’ containing constructs P301S tauRD (cyan), P301S tauRD_QK (sky blue), and P301S tauRD_EL_QK (orange). Data is shown as intensity mean and standard deviation, significance calculated from column effects using 2way ANOVA (*=p<0.05, **=p<0.01, ***=p<0.001, ****=p<0.0001). **b.** Microtubule polymerization assay t_1/2max_ fit to each polymerization curve plotted as an average and standard deviation, significance calculated as in a. **c.** Energetics of residue 301 computed across 2500 MT:tau models for six constructs, colored as in a. The data is shown as violin plots with median and 25/75% quartiles labeled in dashed lines. **d.** Residue energy for tau 275-331 from MT:tau modeled complex for designed constructs. Energies are shown as the mean REU of 2,000 structures colored from high to low energy (yellow to navy)—fragment sequence below with boxed residues signifying the 4R tauRD sequence. Labels above the heatmap denote MT:MAP anchor motifs. Red arrows denote position 301, and blank arrows denote positions 297 and 300, which have lower energy for designed mutations. **e.** Lowest energy structure of modeled sequence for WT tau275-331 (blue, cartoon) and P301S tau275-331 (orange, cartoon) in complex with MT (gray, surface). Black labels denoting MT:MAP anchor points, position 301 labeled for WT pointing down (blue, sticks), and P301S pointing up (orange, sticks).

We similarly tested the 3R tauRD constructs with the polymerization assay at 10 µM and observed similar behavior. Proline to serine constructs had low polymerization, while the 3R-tauRD_DI_VS had deceased polymerization relative to the 3R WT tauRD construct (Supplementary Fig. 7c). Our data recapitulate the observation that the P301S mutations significantly alter MT binding. In the context of the ‘PGGG’ turns, the R1 sequence in the tauRD_EL_QK and 3R WT tauRD appears to increase the polymerization extent without changing the polymerization rate.

### Modeling the MT: tau complex reveals key stabilizing interactions

Local structure in tau has been implicated to be important for its binding to MTs, with the addition of the P301S mutation resulting in an almost complete loss of binding to tau^50^. Available structural models of tau in the context of microtubules include tau in a fully extended conformation determined by cryo-EM^51^ and a more compacted conformation visualized by NMR^52^. To explore how these turn sequence changes may affect the bound state of the MT: tau complex, we modeled the interaction energetics of different tau constructs bound to MTs using an available cryo-EM structure^51^. First, we built WT and mutant 4R tauRD using a threading protocol into the 4xR2 tau from the cryo-EM model (PDB ID 6CVN) to rebuild the missing sequence (residues 300-303), then trimmed the tau model to contain only repeats 2 and 3 (residues 275-331) and minimized the resulting low energy structure in the context of tubulin 1/1/1 trimer. The energetics and conformational heterogeneity were compared for all 6 constructs and we visualized the 10 lowest energy complexes (Supplementary Fig. 7d-f). We compared the energetics of our models by scoring the 301 positions of our MT: tau complex and observed a difference in energetics between the ‘PGGG’ and ‘SGGG’ models (Fig. 5c).

Indeed, plotting the per residue energy contributions of the fragment, we find that the proline to serine yields the most dramatic shift (Fig. 5d, red arrow) and sits firmly between two well-established tubulin binding motifs (Fig. 5d, LXXVXSK motifs)^52,53^. The changes in the energy terms that dictate this difference are attractive and solvation energies suggestive of positive contributions to the burial of the proline ring (Supplementary Fig. 7g). While the serine has the potential to form an additional hydrogen bond, the energy contribution from that interaction does not overcome the increased van der Waals forces (Supplementary Fig. 7g). At the same time, the serine residue points away from the MT (Fig. 5e). Interestingly, we also find that valine to glutamine mutations at position 300 (from tauRD_QK), as well as isoleucine to leucine mutations at position 297 (from tauRD_EL_QK), have lower energy and contribute to complex stability (Fig. 5e, black arrows). This boost in stabilization may explain why the tauRD_EL_QK may yield higher MT binding compared to the WT tauRD (Fig. 5a). By modeling the ‘PGGG’ loop within the available MT: tau structure, there is clear importance of the proline at position 301, suggesting that the proline position not only affect the propensity to aggregate but impact the MT bound state energetics.

### Stabilizing mutations render tau less sensitive to exogenous seeds in cells

Finally, to evaluate whether our designed tau sequences influence response to pathogenic seeds, we developed a cellular assay leveraging mEOS3.2 fusions of tauRD harboring our designed mutants in the context of the pathogenic P301S mutation. To assess the behavior of these mutants, we employed a HEK293T cell model of tauRD aggregation leveraging the FRET-compatible photoconvertible mEOS3.2 system. We produced cell lines stably expressing P301S, P301S tauRD_QK, and P301S tauRD_EL_QK but also single mutants P301S tauRD encoding V300Q (herein P301S tauRD_Q) and P301S tauRD encoding S305K (herein P301S tauRD_K), as fusions to mEOS3.2, which were sorted to maintain similar high expression levels across cell lines. The cells were seeded with a concentration series of recombinant tau fibrils, photoconverted, and then analyzed by flow cytometry (Fig. 6a). To ensure optimal FRET efficiency, the photoconversion time was determined empirically (Supplementary Fig. 8a). Then, we employed a gating strategy to observe the proportion of cells that were FRET-positive (Supplementary Fig 8b). As observed previously, P301S tauRD-mEOS3.2 yielded high percentages of cells, with 74±4% of FRET-positive cells seeded with 100 nM recombinant tau fibrils (Fig. 6b, green). The introduction of V300Q and S305K mutations alone reduced seeding 1.2-fold (Fig. 6b, yellow and orange) while combining these mutations in the P301S tauRD_QK reduced seeding activity nearly 2-fold (Fig. 6b, blue). Finally, introducing the P301S tauRD_EL_QK reduces seeding activity by 2-fold compared to the P301S tauRD control (Fig. 6b, red). Representative images of cells expressing each of the 6 tauRD constructs controlled or seeded with 100 nM recombinant tau fibrils show a reduction in puncta frequency (Fig. 6c). This data reveals that modification of the turn residues V300Q and S305K is sufficient to reduce P301S tauRD sensitivity to pathogenic tau seeds in cells.

**Figure 6.**
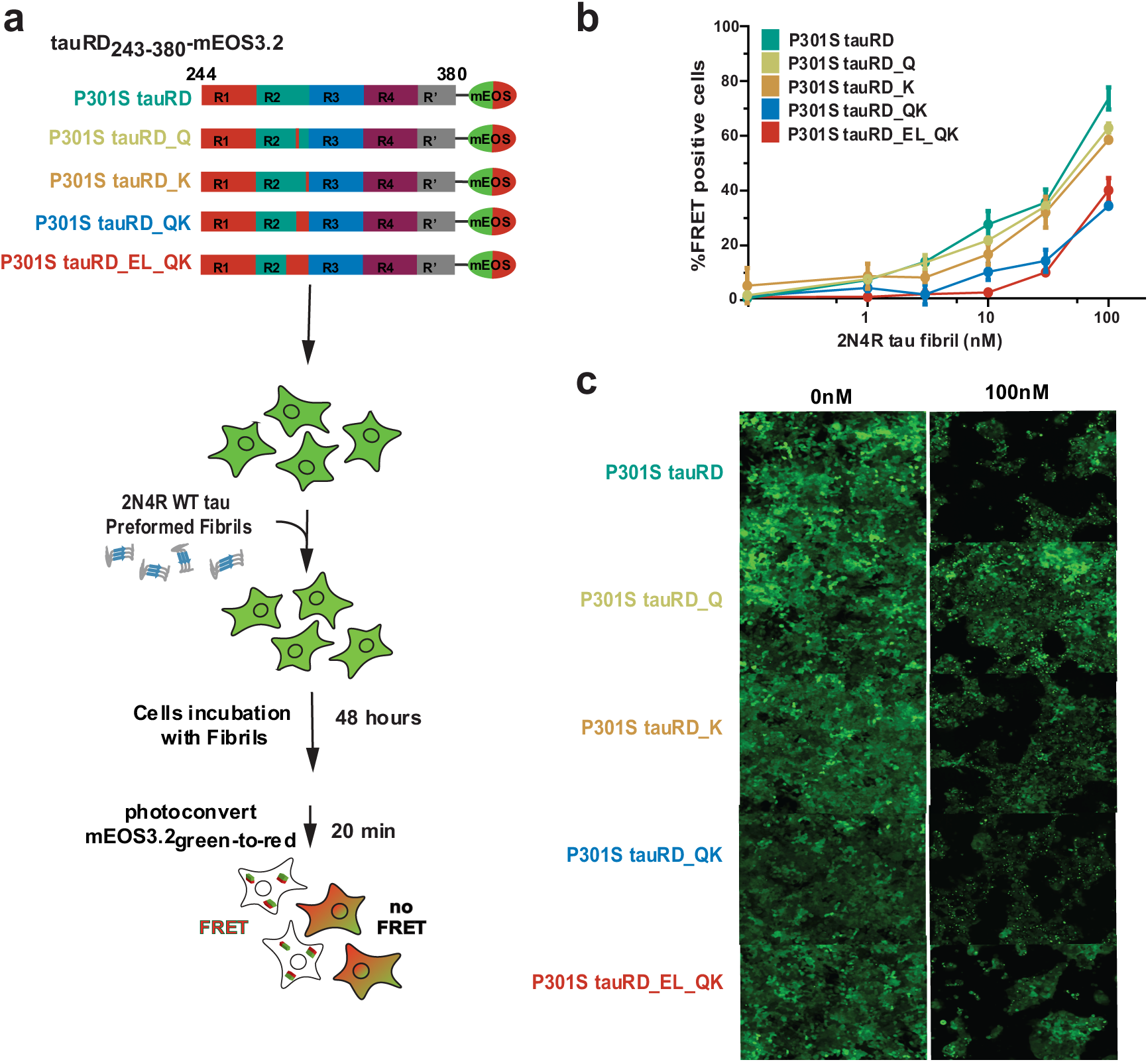
Designed tau sequences reduce tau aggregation in cells. **a.** Schematic of fibril propagation seeding workflow. Cartoon of 4R tauRD P301S designed constructs encoding 3R-like mutants as fusions to mEOS3.2, used for lentiviral transfection into HEK 293T cell and sorted for high expression, treated with lipofectamine mediated seeding from 2N4R WT tau preformed fibrils, followed by incubation time, photoconversion and analysis by percent FRET positivity. **b.** Percent FRET positivity (FRET pairs: tauRD-mEOS3.2green/tauRD mEOS3.2red) was measured by flow cytometry on n = 3 biological replicates of at least 10,000 cells per fibril concentrations and expression construct. Data are shown as averages with standard deviation. **c.** Imaging of HEK 293T cells expressing designed constructs 48hrs after no treatment control and max treatment of 100 nM fibril concentration.

## DISCUSSION

The conversion of tau into insoluble aggregates is linked to several neurodegenerative diseases. Tau isoform conversion between 3R and 4R leads to different aggregation propensities; nevertheless, minor sequence changes surrounding amyloidogenic motifs provide mechanistic hints toward the intrinsic regulation of tau aggregation. We find that the conserved proline in the ‘PGGG’ motif, coupled with the residues that flank it, plays a vital role in limiting the self-assembly of the amyloidogenic ^306^VQIVYK^311^. This region adopts multiple conformations in the end-state of the tau fibril fold, providing multiple residues that can form stabilizing interactions with the amyloid motif. We observe that the local rearrangements driven by P301S mediate the exposure of the amyloid motif in the 4R context, which the 3R-based designed sequence can reverse. Introducing the P301S mutation in the 4R-derived fragment yields more disordered and dynamic states for glycine 302, reducing local and long-range interactions and increasing the exposure of the amyloid motif. This P301S destabilizing effect is minimized in the context of only two residue swaps, V300Q and S305K, with minimal changes in the dynamics. With these mutations, we see a rearrangement at the ‘PGGG’/’SGGG’ region, increasing intermediate-range interactions. We propose a structural model for how the proline to serine mutation is the main driver of alterations in binding to MTs. Yet, we identify differential interactions for the 294-300 residues to MTs between 3R and 4R tau that increased microtubule polymerization when containing a ‘PGGG’ motif. We finally detect decreased sensitivity to pathogenic seeds through our paired mutations. Together, our data suggest an avenue for tuning tau aggregation through small local structure rearrangements that alter turn dynamics which retain biological activity.

FTD-linked pathogenic mutations largely localize to the repeat domain of MAPT and, more specifically, cluster at inter-repeat elements^54^, including specific mutations in the variable exon 10 of *MAPT* associated with altering splicing^54^. A unifying mechanism for how a diversity of FTD-linked mutations drive the same disease is unclear; cryo-em structures of tau amyloids isolated from FTD-tau patients may provide hints in the future. It remains unknown how these mutations lead to tau-based dysfunction. Still, the specificity of the pattern of mutations in MAPT may suggest that loss-of-function (LOF) and gain-of-function (GOF) mechanisms contribute to pathological fibrilization and impaired microtubule binding, ultimately leading to FTD-tau. The mutation sites often localize in proximity to amyloid motifs, suggesting that mutations may disrupt biological function and promote amyloid-motif-dependent tau self-assembly. Indeed, analyzing disease-associated mutations in extensive repositories of globular proteins (using AlphaFold models) reveals distinct patterns: LOF mutations are randomly distributed across the protein structure. In contrast, GOF mutations cluster at sites critical to biological function, where they cause disruption^15^. Applying this concept to MAPT may hint at a unifying mechanism of FTD-tau disease. Developing new strategies to modulate tau is essential to better understand how to mitigate GOF and LOF effects.

The field has focused on understanding the pathological effects of endpoint aggregates to guide potential ways to diagnose or treat disease. How tau interacts with MTs has garnered less interest. It remains unclear what features of microtubule-associated proteins (MAPs) are required for binding to MTs. The *MAPT* gene is highly conserved across evolution^53^. Similarly, the related *MAP2* and *MAP4* genes are conserved but do not encode aggregation-prone sequences^53^. These genes are processed similarly during splicing, producing both 3R and 4R isoforms of the sequence that retain binding to MTs but are not implicated in misfolding diseases. Two amino acids in the tau amyloid motif, **V**Q**I**VYK, are conserved across MAPs. Because this site is essential for splicing, and even codon changes can alter splicing, part of the amyloid motif may be preserved to retain efficient splicing. However, it is still being determined why the latter part of the amyloid motif is conserved. Our data demonstrate that substituting two residues flanking the turn preceding the VQIVYK motif in tau to mimic the 3R localized sequence reduces 4R tau aggregation and maintains tau’s ability to stabilize MTs. Prior cryo-EM structural data on tau bound to MTs showed a fully extended conformation, highly disordered at the inter-repeat regions surrounding the ‘PGGG’ motif. By contrast, NMR studies showed a relatively collapsed conformation of this element bound to MTs, adopting a turn conformation around the ‘PGGG’ motif. Our tau: MT models of the bound state suggest a role for the wild-type proline the interaction complex.

Future experiments and analysis must more closely interrogate the relationship between the tau monomer ensemble in solution, MT binding states, and the early misfolding steps that lead to disease. We suspect these intermediate and long-range interactions may be important in forming the MT: tau encounter complex. Interestingly, XL-MS experiments on tauRD encoding mutations that convert the local sequence from 4R-like to 3R-like alter the frequency of global long-range contacts but increase intermediate-range contacts. This shift in frequency of interactions correlates with stabilization of the ‘PGGG’ turn to reduce aggregation and maintain or even increase MT activity. Future experiments may explore how the computational design of tau may lead to mutants that can replace wild-type MAPT to reduce disease. A deeper comparison of naturally occurring MAP sequences may offer clues for the design of tau sequences that retain MT binding but aggregate less. Thus, more focus must be brought onto the effect of mutations on the multi-state problem of tau conformation in solution (aggregation-resistant), tau bound to MT (functional state), and tau misfolding towards pathological aggregates to begin to understand the origins of tau in function and disease.

A deeper interpretation of the evolvability of tau to tune these two properties may be inferred from the evolutionary sequence history of MAPs. Future studies must test the biological consequences of designer MAPT sequences in animal models as ongoing therapeutic approaches to reduce tau levels may lead to unknown consequences in humans.

## METHODS

### Peptide synthesis and disaggregation

All peptides were synthesized as ordered by Genscript with N-terminal acetylation and C-terminal amidation modifications at >95% purity. Peptides were disaggregated as previously described^55^. In brief, peptides were resuspended in a 250 µL Trifluoroacetic acid (TFA) (Pierce) incubated at room temperature (RT) for 1-16 hours in a chemical fume hood. The peptide solution was dried under a stream of nitrogen gas, and then 100 µL of H_2_O was added, followed by flash freezing, and immediately placed under vacuum to remove any residual solvents. The peptide residue was resuspended in 200 µL of H_2_O to determine the concentration and, with 10X PBS pH 7.4, adjusted with NaOH and H_2_O to achieve the desired pH and concentration.

### ThT fluorescence aggregation assays

Peptide concentrations were adjusted with 1X PBS (136.5 mM NaCl, 2.7 mM KCl, 10 mM Na_2_HPO_4_, 1.8 mM KH_2_PO_4_, pH 7.4) and 1 mM Dithiothreitol (DTT) to a desired concentration between 800 µM and 50 µM. Thioflavin T (ThT) was added to the samples at a final concentration of 25 µM. Wild-type or mutant 4R or 3R tauRD protein was diluted in 1X PBS with 10 mM DTT to the desired concentration of 40, 20, 10, and 5 µM. A 2X heparin concentration (Amsbio) was added to the tauRD protein. All samples were aliquoted at 45 µL per replicate in a 384-well clear bottom plate. All conditions were performed as three independent replicates at 37 1C. ThT kinetic scans were run every 30 min, with 10 seconds of orbital shaking before each read, on a Tecan Spark plate reader at 446 nm Ex (10 nm bandwidth) and 482 nm Em (10 nm bandwidth). Blank wells containing buffer and ThT were subtracted from experimental values. The data were plotted, and the *t*_1/2_ values were estimated using a non-linear regression model fitting in GraphPad Prism and reported as averages with a standard deviation.

### Peptide NMR and secondary structure calculation from chemical shifts

NMR spectra were acquired on Agilent DD2 spectrometers operating at 600 MHz at the UTSW Biomolecular NMR Facility. ^1^H-^1^H TOCSY spectra^56^ and ^1^H-^1^H NOESY spectra^56^ were acquired at 25 1C. Peptides were adjusted to 500 µM in 1X PBS, 1mM DTT with 10% D_2_O or 99.998% D_2_O, and the total acquisition time for ^1^H-^1^H TOCSY and ^1^H-^1^H NOESY spectra ranged from 8-11 hours. Five ^1^H-^1^H TOCSY spectra were collected in series for the aggregation-prone P301S R2R3_294-311_ peptide, each 10 hours long. All data were processed with NMRpipe^57^ and analyzed with CARA (http://cara.nmr.ch/doku.php) and SPARKY^58^. Secondary structure propensity was calculated using the neighbor-corrected secondary structure propensity calculator (ncSPC) from the Ha chemical shift assignments. The calculator used the deviations from random coil chemical shifts to estimate the proportion of secondary structure per residue corrected by a specified neighbor window. For our analysis, we used a neighboring residue window of 3^36,37^. The data were plotted in GraphPad Prism.

### Tau amyloid fibril structure comparison

Evaluated fibril structures were taken from previous analyses of structural stability (PDB accession codes: 5o3l, 5o3t, 6npw, 7p66, 6tjo, 7p6d, 7p6a, 7p65, 6gx5, 8q92). Folds categorized as described Alzheimer’s disease (AD), chronic traumatic encephalopathy (CTE), Globular glial tauopathy (GGT), corticobasal degeneration (CBD), Argyrophilic grain disease (AGD), Progressive supranuclear palsy (PSP), GGT/PSP-fold (GPT), Picks disease (PiD), and PS19 AD mouse model with human tau derived fibrils (P301Sm). All structures were aligned based on Cß positions of the ^295^VQIVYK^311^ motif and images created using Pymol. Solvent accessible surface area (SASA) was calculated for the middle chain of the fibril. Then, the per residue normalized SASA value was calculated as a ratio to the maximum estimated SASA of each residue as described in Tien et al. 2013^59^—normalized DDG values obtained from Mullapudi et al. 2023{Mullapudi, 2023 #4}. New values were calculated for P301Sm fibril (PDB accession code 8q92) following the previously described protocol {Mullapudi, 2023 #4}. Briefly, the PDB structure was expanded to obtain 9 layers of tau monomer. The flex-DDG fibril alanine scan method was implemented, and the change in ΔG between the alanine mutant and the WT structure at each position was calculated. All DDG values were min-max normalized to each structure’s 295-311 region.

### Molecular dynamics simulations production

Molecular Dynamics simulations were performed using PLUMED patched GROMACS v2018.4. Each peptide was simulated in a dodecahedron box with SPC/E water starting from a fully extended conformation built-in Pymol for a total of 5 μs followed by 15 μs in TIP3 water with 150 mM NaCl ions. The initial dodecahedron simulation box was constructed using the extended peptide structure with a minimum 1.5 nm boundary condition. The AMBER99sb-ildn forcefield^60^ was used for all simulations. After an initial 1000 steepest descent steps of converged energy minimization, 10 ns of NVT and 20 ns of NPT (first 10 with Berendsen^61^ and the last 10 with Parrinello-Rahman^62^ barostats) equilibrations were performed.

The subsequent production level trajectories are based on 5 fs time steps using hydrogen-only virtual sites^63^. Production level trajectories were obtained for an NPT ensemble with Parrinello-Rahman barostat and periodic boundary conditions with Particle Mesh Ewald (PME)^64^ summation for long-range electrostatics. A total of 20 μs trajectories were generated by producing five 1 μs simulations per peptide. These trajectories were analyzed using the GROMACS clustering function with a cluster cutoff of 0.54 nm. The cluster cutoff was chosen empirically to produce a maximum of 15 clusters for all four peptides. The center structure of the top 5 clusters was then used as the starting structure for simulations with Na+ and Cl- ions included in the solvent. These second-stage simulations were performed with the same minimization, equilibration, and production workflow with only a few modifications. A dodecahedron simulation box was used, using the new starting structures with an increase in the minimum boundary cutoff to 1.6 nm. The same AMBER99sb-ildn forcefield^60^ was used with an update to a TIP3P water model with 150 mM ionic strength by including Na+ and Cl- ions. The energy minimization step was performed with the steepest decent algorithm to obtain a maximum force <1000.0 kJ/mol/nm. Equilibration was run aligned with the previous step. The subsequent production level trajectories were performed as the initial simulations, with an updated 2 fs timestep to account for the ionic solvent. A total of 60 μs trajectories were generated by producing five 3 μs trajectories per peptide. All simulations were done on UTSW’s BioHPC computing cluster.

### MD analysis: clustering, RMSD, and distance calculations

All analysis was done using GROMACS 5.0.4^70^. The analysis framework was done as previously described^55^. Specifically, backbone distance in Fig. 2b and Supplemental Fig. 3a was calculated using the ‘gmx pairdist’ command between the center of mass of backbone atoms for residues 1-6 (i.e., XXNXKH) vs. residues 12-18 (i.e., XVQIVYK) from the 15 μs ensemble of the second production stage and the 5 μs of the initial replicate simulations. Data is shown as violin plots (e.g., density distributions) with mean and 25% and 75% quartile marked. For the second step of the simulation, the 5 independent trajectories were concatenated into a 15 μs ensemble, and the 2-dimensional RMSD was calculated every 10 ns using the ‘gmx rms’ command. The 15 µs ensembles were also compared to the published structure of tau. The RMSD of the ensemble referenced the P301S fibril structure (PDB ID 8Q96), which was reported as a distribution violin plot in Supplemental Fig. 3e. All graphs were plotted directly from gmx output or using GraphPad Prism. Images were created using Pymol for cluster structures.

### MD state analysis with Markov State Model (MSM)

MSM of the peptide simulations was built from the combined ensemble of all four peptides and the individual peptide ensembles, concatenating each peptide 15 μs simulations in ionic solvent using the pyEMMA software. We constructed the MSM of these trajectories to capture the kinetic behavior of the system with the PyEMMA 2.5.12 library^40^ in Python 3.10.9. After the second step of MD simulation WT R1R3_263-311ΔR2_, P270S R1R3_263-311ΔR2_, WT R2R3_295-311_, and P301S R2R3_295-311_, we trimmed the 50 μs trajectory at 100 ps timestep, and each resulting trajectory (5 concatenated trajectories for 4 systems) contains 30001 frames. These trajectories were trimmed again to contain only Cα atoms and concatenated to generate a 60,000,001-frame trajectory. The MSM featurization step was performed based on the minimum pairwise distance between residues 1-7 (i.e., TENLKHQ vs. KDNIKHV) against amyloid motif residues 12-18 (i.e., XVQIVYK) with a total 49 distance pairs with 150001 frames for a single system. We considered the four featurization techniques for model construction: torsion angle, backbone atom positions, backbone atom intra-molecular distances, and minimum pairwise distance between amyloid motifs. The featurization criteria were chosen based on VAMP-2 scores (Variational Approach for Markov Process).

We selected the minimum pairwise residue distance over the torsion angles because these motifs covered functional conformational change and reported on interactions across the peptide region that differ in sequence. Further dimensionality reduction was performed from a total of 49 pair distances (600000001,49; 150001,49) to 2 dimensions (600000001,49; 150001,2) using the time-lagged Independent Component Analysis (tICA). TICA identifies the slowest mode of feature space and maximizes the autocorrelation of reduced coordinates for state decomposition. After that, we used the k-means clustering algorithm for the decomposition of reduced conformational space into 200 microstates. Those conformations that belong to the same energy basin in the TICA subspace are clustered into each microstate, and transition occurs quickly among these states. Further, we lumped the microstates into 6 macrostates using PCCA+ to predict the major conformational changes in peptides. The MSM was determined on an implied time scale plot, and the lag time 200 (time/step) was chosen. The Chapman-Kolmogorov test was applied to validate the consistency of probabilities predicted from MSM and MD simulations. The calculated MSM was further coarse-grained into hidden Markov models to calculate the metastable macrostates and transition probabilities. Two hundred representative structures were computed for each state extracted from the centroid of the most populated microstate. We calculated the pairwise distance for all residue pairs for the 200 structures of each cluster using ‘gmx pairdist’ in gromacs and plotted the mean distance and SD in heatmaps using MATLAB plotting software. The first 10 structures were used as representative models for each state. Images were created using PyEMMA graphics output options or Pymol for representative structures.

### Recombinant tauRD and full-length tau production

We utilized several forms of recombinant tau. Recombinant repeat domain (tauRD) 4R isoform (4R tauRD, 4R P301S tauRD, 4R tauRD_EL_QK, 4R tauRD_EL_QK, and 4R P301S tauRD_QK) and 3R isoform (3R tauRD, 3R P270S tauRD, 3R tauRD tauRD_VS_DI and 3R P301S tauRD tauRD_VS_DI) constructs were purchased from Twist Bioscience encoded into pET-29b(+) vector without tags. Each plasmid was transformed into BL21-Gold (DE3) cells. Cells were grown in 1X Terrific Broth media with 1 μg/ml kanamycin to OD600 of 1.4 and induced with 1 mM isopropyl β-D-1-thiogalactopyranoside (IPTG) for 3 h at 37 °C. To produce N ^15^ isotope-labeled samples of 4R tauRD, 4R tauRD P301S, 4R tauRD_EL_QK, and 4R tauRD P301S tauRD_EL_QK, a single colony of freshly transformed plasmid into BL21(DE3) was inoculated into 10mL 2xLB supplemented with 0.05mg/mL Kanamycin and incubated for 7-8 hours at 37°C while shaking at 220rpm. The 10mL culture was then mixed into 100mL of M9 minimal media (42mM Na_2_HPO_4_, 22mM KH_2_PO_4_, 8.5mM NaCl, 0.1mM CaCl, 2mM Mg_2_SO_4_, 1E-4% Thiamine, 0.4% Glucose, 187mM NH_4_Cl, 0.05mg/mL Kanamycin) and incubated overnight at 37°C while shaking at 220rpm. In the morning of the following day, the cells were spun down at 2,000g for 10 minutes and resuspended in 20 mL of M9 minimal media containing 15N labeled NH_4_Cl in place of the unlabeled molecule. This resuspension was immediately added to 1L of M9 minimal media with ^15^N labeled NH_4_Cl and incubated at 37°C shaking at 220rpm. Once OD600 reached 0.6-0.8 A.U., 1mL of 1M IPTG was added to induce protein expression for 3 hrs at 37°C. The cells were harvested and lysed in 20 mM 2-(N-morpholino) ethane sulfonic acid (MES), 1 mM Ethylenediaminetetraacetic acid (EDTA), 1 mM MgCl, 5 mM β-mercaptoethanol (BME), 1 mM PMSF, pH 6.8, using an Omni Sonic Ruptor 400 with lysate on ice. After sonication, 500 mM NaCl was added to the lysates, which were boiled for 20 minutes before clarifying by centrifugation. The supernatant of the clarified lysate was exchanged into 20 mM MES, 50 mM NaCl, 5 mM BME, and pH 6.8 by dialysis. The dialyzed sample was filtered, applied to a HiTrap SP HP (GE), and eluted with a 50 mM–1 M NaCl gradient. Tau-containing fractions were concentrated on an Amicon-15 concentrator (5KD cutoff) and applied to HiLoad 16/600 Superdex 75 pg (GE) and eluted into 1X PBS (136.5 mM NaCl, 2.7 mM KCl, 10 mM Na_2_HPO_4_, 1.8 mM KH_2_PO_4_, pH 7.4), 10 mM DTT. Aliquots were all stored at −80 °C in 1 X PBS.

The pET-28b-tau plasmid encoding full-length 2N4R WT tau was a gift from Dr. David Eisenberg (UCLA). The plasmid was transformed into BL21-Gold (DE3) cells. Cells were grown in 1X Terrific Broth media to OD600 of 1.4 and induced with 1 mM isopropyl β-D-1-thiogalactopyranoside for 3 h at 37 °C. The cells were harvested and lysed in 50 mM Tris, 500 mM NaCl, 1 mM β-mercaptoethanol, 20 mM imidazole, 1 mM phenylmethylsulfonyl fluoride (PMSF), pH 7.5, using an Omni Sonic Ruptor 400 at 4 °C. The lysates were centrifuged at 13,000 x g for 20 minutes at 4 °C, and the supernatant was applied to a Ni-NTA gravity column and eluted with 50 mM Tris, 250 mM NaCl, 1 mM β-mercaptoethanol, and 300 mM imidazole. Eluting fractions containing tau were desalted into 50 mM MES, 50 mM NaCl, and 1 mM β-mercaptoethanol (pH 6.0) by PD-10 column (GE). Exchanged fractions were applied to a HiTrap SP HP (GE) and eluted with a 50 mM–1 M NaCl gradient. Tau-containing fractions were concentrated on an Amicon-15 concentrator (10kDa cutoff) and applied to a HiLoad 16/600 Superdex 200 pg (GE) and eluted into 1X PBS (136.5 mM NaCl, 2.7 mM KCl, 10 mM Na_2_HPO_4_, 1.8 mM KH_2_PO_4_, pH 7.4). Aliquots were all stored at −80 °C in 1X PBS.

### 4R tauRD NMR experiments

^1^H-^15^N HSQC spectra, ^1^H-^15^N NOESY-HSQC, and ^1^H-^15^N TOCSY-HSQC spectra were acquired at 10 1C. NMR spectra were acquired on Agilent DD2 spectrometers operating at 800 MHz at the UTSW Biomolecular NMR Facility. tauRD constructs were concentrated to 200-300µM in tauRD NMR buffer (10mM phosphate buffer, 100mM NaCl, and 4mM DTT with 10% D_2_O), replicating buffer conditions of previously published data. The concentration of tauRD constructs was estimated using 215nm absorbance and verified to have a comparable signal-to-noise for NMR spectra. The total acquisition time ranged from 64-80 hours for ^1^H-^15^N NOESY-HSQC and ^1^H-^15^N TOCSY-HSQC spectra. The four tauRD constructs were then diluted to 50uM, and ^1^H-^15^N HSQC spectra were measured for 4 hours. All data was processed with NMRpipe^57^ and analyzed with CARA (http://cara.nmr.ch/doku.php), SPARKY^58^, and ccpNMR^65^. Peak assignments were transferred from deposited data for tau K18 (bmrb 19253)^66^, and additional N-termini peaks were assigned from htau4p (bmrb 50701)^67^. Mutations in tauRD_EL_QK and P301S tauRD_EL_QK were assigned based on sidechain assignments in ^1^H-^15^N NOESY-HSQC and ^1^H-^15^N TOCSY-HSQC and assignments of K19 tau made from 3R tauRD fragment^68^ (Visualized Source Data 6). No assignments were made for the V300Q mutation.

(^1^H)-^15^N heteronuclear NOE relaxation data (hetNOE) for 150-300 μM of 4R tauRD, P301S tauRD, tauRD_EL_QK, and P301S tauRD_EL_QK samples in tauRD NMR buffer 10mM were obtained using standard Bruker pulse sequences acquired at 10 °C in an 800MHz (^1^H) Bruker AVNEO spectrometer equipped with a Z-gradient cryoprobe. Protein concentrations were estimated using a 215nm absorbance signal; as stated previously, peak intensities were compared across samples to ensure a 20% difference in intensities. Similar experiments were performed at 25 °C for 300uM P301S tauRD and P301S tauRD_EL_QK, and transferring assignments was not possible as published data is at 10 °C and peak intensities were lost or shifted for most peaks, interpreting the data impossible. Limited assigned peaks and intensity differences are available in Source Data 5. In all relaxation experiments, the spectral width was 7812.5 Hz for ^1^H and 1824.25 Hz for ^15^N dimensions. HetNOEs were calculated from the ratio of cross-peak intensities in spectra collected with and without amide proton saturation during the recycle delay. Uncertainties in peak heights were determined from the standard deviation of the intensity distribution in signal-less spectral regions. All NMR spectra were processed in Topspin 4.1.1 and analyzed in Sparky 3.1969^58^. The data were plotted in GraphPad Prism.

### Transmission electron microscopy

An aliquot of 5 μL of the sample was placed onto a glow-discharged Carbon/Formvar-coated 400-mesh copper grid for 1 minute, washed with distilled water, and then negatively stained with 2 μL of 2% uranyl acetate for 1 minute. Images were acquired on a Tecnai G^2^ spirit transmission electron microscope (FEI, Hillsboro, OR), serial number D1067, equipped with a LaB_6_ source at 120 kV using a Gatan ultrascan CCD camera.

### Gel-extracted sample preparation for cross-linking mass spectrometry analysis

174 µg of each construct of 4R tauRD (WT, QK, EL_QK, P301S, P301S QK, and P301S_EL_QK) was diluted in 1X PBS, 10 mM DTT to a final concentration of 58 µM. The cross-linking reaction was conducted by adding disuccinimidyl suberate (DSS, DSS-d0 and -d12, Creative Molecules, dissolved in DMF) to a final concentration of 1 mM for three minutes at 37 °C while shaking at 350 RPM. The reaction was then quenched by adding ammonium bicarbonate to 50 mM final concentration and incubating for 30 minutes at 37 °C while shaking at 350 RPM. Cross-linked samples were resolved on SDS-PAGE gels (NuPAGE™, 4 to 12%, Bis-Tris, 1.5 mm), and the monomer band was extracted from the gel for XL-MS analysis using the following protocol. Bands were sliced into 1mm^3^ pieces and washed twice with MQ water, discarding supernatant after each time. Gel pieces were then covered in acetonitrile/50 mM ammonium bicarbonate (mixed in ratio 2:3, v/v) and sonicated for 5 minutes, and the supernatants were discarded each time. This step was repeated twice. Next, gel pieces were washed in 100% acetonitrile until they became white, then the liquid was removed, and gel pieces were dried by lyophilization. Subsequently, gel pieces were covered in 25 mM ammonium bicarbonate with 10 mM DTT and incubated for 1 hour at 56 °C while shaking at 500 RPM. Next, samples were cooled down, supernatants removed, and a freshly prepared 25 mM ammonium bicarbonate solution with 55 mM iodoacetamide was added. Gel pieces were incubated for 45 minutes in the dark at room temperature. Then, gel pieces were sequentially washed with 25 mM ammonium bicarbonate, 50% acetonitrile, and 100% acetonitrile with quick vertexing after each wash. The liquid was removed, and gel pieces were dried by lyophilization. Trypsin digestion was initiated by adding 1:10 (trypsin to protein, m/m) mass spectrometry grade trypsin (New England Biolabs #P8101S) in 50 mM ammonium bicarbonate. Then, the reaction was incubated overnight at 37 °C while shaking at 500 RPM. Digestion was stopped by adding 2% (v/v) formic acid, and supernatants were collected. Peptides were additionally extracted by covering gel pieces in MQ water/acetonitrile/formic acid (50:50:0.1, v/v/v), incubated for 15 minutes at 37 °C while shaking at 500 RPM, and then sonicated for 5 minutes. This step was repeated twice, and collected supernatants were pooled, snap-frozen, and lyophilized. Peptides were resuspended in 50 mM ammonium bicarbonate/acetonitrile/formic acid (93:5:2), further purified by solid phase extraction using Sep-Pak tC18 cartridges (Waters #WAT054960), snap frozen and lyophilized. Samples were processed as 5 technical replicates on a Thermo Orbitrap Fusion Lumos at UTSW Proteomics core. The mass spectrometer was operated in data-dependent mode by selecting the five most abundant precursor ions (m/z 350–1600, charge state 3+ and above) from a preview scan and subjecting them to collision-induced dissociation (normalized collision energy = 35%, 30 ms activation). Fragment ions were detected at low resolution in the linear ion trap. Dynamic exclusion was enabled (repeat count 1, exclusion duration 30 s). Data was processed to identify unmodified, 1 mono-link, 2 mono-links, and loop-links by peak areas. Data was reported as fraction modification normalized to the total peak area per replicate.

### Analysis of mass spectrometry data

Thermo.raw files were converted to the open.mzXML format using msconvert (proteowizard.sourceforge.net) and analyzed using an in-house version of xQuest^46^. Spectral pairs with a precursor mass difference of 12.075321 Da were extracted and searched against the respective FASTA databases containing tau (TAU_HUMAN P10636-8) or designed constructs with and without P301S mutations. xQuest settings were as follows: Maximum number of missed cleavages (excluding the cross-linking site) = 2, peptide length = 5–50 aa, fixed modifications = carbamidomethyl-Cys (mass shift = 57.021460 Da), variable modification = oxidation of methionine (mass shift = 15.99491 Da), mass shift of the light crosslinker = 138.0680796 Da, mass shift of mono-links = 156.0786442 and 155.0964278 Da, MS1 tolerance = 10 ppm, MS2 tolerance = 0.2 Da for common ions and 0.3 Da for cross-link ions, search in ion-tag mode. Post-search manual validation and filtering were performed using the following criteria: xQuest score >25, FDR <0.05, mass error between − 2.2 and + 3.8 ppm, %TIC >10, and a minimum peptide length of six aa. FDRs for the identified cross-links were estimated using xProphet^69^. The five replicate data sets were compared, and cross-links in five out of five data sets were used to generate a consensus data set. Normalized mono-links and cross-link data with information on cross-linked residue positions and nseen (frequency) were visualized using customized Matlab scripts. Hierarchical clustering was done using Matlab tools clustergram and linkage to build linkage dendrogram trees across identified loop-links/cross-links for all 6 constructs.

### Tau aggregation in cells using mEOS biosensor cell lines

FM5-CMV constructs of 4R P301S, P301S_Q, P301S_K, P301S_QK, and P301S EL_QK fused to mEOS3.2 at the C-terminal were used for cell expression. The virus of each construct was produced in Lenti-X™ 293 T Cell Line (Takara, Cat. #632180) and previously described^70^. Specifically, 300 ng PSP helper plasmid, 100 ng VSVG, and 100 ng of the plasmid of interest were transiently co-transfected using 1.875 µL TransIT (Mirus Bio), and Opti-MEM (Gibco) were mixed to a final volume of ∼30 µL. Media (10% FBS, 1% Pen/Strep, 1% GlutaMax in Dulbecco’s modified Eagle’s medium) containing the virus was collected after 48 h and centrifuged at 1000 RPM for 5 min to remove debris and dead cells. For transduction, 10-50 µL of virus suspension was added to HEK293T (ATCC CRL-1268), and cells were grown in virus-containing media for 72 h before expanding to the most confluent condition. Cells were harvested from a 10 cm dish with 0.05% trypsin, resuspended in flow cytometry buffer (HBSS plus 1% FBS and 1 mM EDTA), and subjected to FACS (Sony Biotechnology). Populations with FITC-A signal above 10^4^ were collected for high-expressing cells. Following FACS and expansion, cells were maintained as a polyclonal line. All stable cell lines were amplified, frozen, and stored in liquid nitrogen until use.

For seeding experiments, cells were plated in 96-well plates at 20,000 cells/well in 130 µL of media. 24 h later, the cells were treated with 20 µL of fibril treatment with a dilution series in technical triplicates. Briefly, treatments were composed of heparin-induced 2N4R recombinant tau fibrils (8 µM monomer equivalent), sonicated for 5 minutes at an amplitude of 65 and 30 s off/on a Q700 Sonicator (QSonica). Sonicated fibrils were diluted to the desired concentration at 10 µL, mixed with 1 µL lipofectamine and 9 µL Opti-MEM, incubated at RT for 30 min per condition, and replicated. After incubation, samples were added to cell media for final concentration in the well of 300 nM, 100 nM, 30 nM, 10 nM, 3 nM, 1 nM, and media control. After 48 hours, cells were harvested with 0.05% trypsin and then fixed in 2% paraformaldehyde (Electron Microscopy Services) for 10 min at room temperature. PFA was removed, and cells were resuspended in 1X PBS. For the initial experiment, the optimal time of photoconversion was determined by exposing cells to UV for 0-60 minutes and measuring the mEOS red-to-green ratio, which determined 20 minutes as the optimal Red/Green ratio used for all subsequent experiments.

### Flow cytometry

A BD LSRFortessa was used to perform FRET flow cytometry. The mCerulean3 and FRET signal was measured by exciting with the 405 nm laser, and fluorescence was captured with a 405/50 nm and 525/50 nm filter, respectively. mClover3 signal was measured by exciting cells with a 488 nm laser, and fluorescence was captured with a 525/50 nm filter. To quantify FRET, we used a gating strategy where mCerulean3 bleed through into the mClover3, and FRET channels were compensated using FlowJo analysis software, as described previously^71^. Subsequently, we created a final bivariate plot of FRET vs. Cerulean and introduced a triangular gate to assess the number of FRET-positive cells. This FRET gate was adjusted to biosensor cells that received lipofectamine alone and are thus FRET-negative. FRET signal is defined as the percentage of FRET-positive cells in all analyses. For each experiment, 10,000 mClover3/mCerulean3 double-positive cells per replicate were analyzed, and each condition was analyzed in triplicate. Data analysis was performed using FlowJo v10 software (Treestar).

For the mEOS3.2 cell expression system, tauRD-mEOS biosensor cells were first photoconverted under UV for 20 min. 10,000 singlet events corresponding to the donor (non-photoconverted mEOS3.2) and acceptor (photoconverted mEOS3.2) positive cells were collected for each sample. Non-photoconverted mEOS3.2 was collected at the Alexa Flour 488 channel. FCS files were exported from the BD FACSDiva data collection software and analyzed using FlowJo v10 software (Treestar). Compensation was manually applied to correct donor bleed-through into the FRET channel guided by a sample with non-aggregated and photoconverted tauRD-mEOS. Samples were gated on the acceptor intensity such that cells with similar concentrations of tauRD-mEOS were analyzed to mitigate the contribution of differences in concentration leading to apparent changes in the fraction of FRET-positive cells in each condition. FRET-positive cells were quantified by gating double-positive singlet events with a ratio of FRET to donor signal higher than that of a population of tauRD-mEOS photoconverted cells without aggregates.

### Microtubule Polymerization assay

Experiments were carried out following the Cytoskeleton Tubulin Polymerization Fluorescence-based Assay (BK011P) protocol. TauRD constructs in 20 mM MES and 10 mM DTT buffer were mixed with 99.9% purified tubulin at 2 mg/ml in 80 mM PIPES, 2 mM MgCl, 0.5 mM EGTA, pH 6.9, 10 µM fluorescent reporter to a final concentration at 10 µM. 3 µM paclitaxel solution was used as a positive control. All conditions were done in triplicates at 37 °C. ThT kinetic scans were run every minute on a Tecan Spark plate reader at 350 nm Ex (10 nm bandwidth) and 435 nm Em (20 nm bandwidth). Blank wells containing buffer and tubulin were subtracted from experimental values. The data were min-max normalized to the sample with maximal signal (Taxol positive control), and the data was used for non-linear regression to determine the intensity max and the t_1/2_max for assessing polymerization time. Column effects using 2way ANOVA were calculated for the intensity max and t_1/2_max using GraphPad Prism v8.0-10.0.

### MT:tau structure Analysis

The synthetic tau (R2X4) model bound to microtubules (MT) from PDB ID 6CVN was rebuilt to have the 4R tauRD designed sequence using the Rosetta threading protocol with bilateral loop building to include the missing residue from the structures^51^. For each sequence, 2,500 models of only tau without the MT were produced, constrained by the initial template of the tau structures. Models were then realigned with the α/β/α tubulin, and the tau chain was trimmed to contain only tau repeats 2 and 3 (275-331) with backbone atoms only (atoms NH, N, CA, CB, and C=O). Each structure was minimized using the Rosetta protocol with the algorithm and a minimum tolerance of 0.001 (https://www.rosettacommons.org/software). The resulting minimized structures were scored as a complex using the InterfaceAnalyzer protocol, pre-packing separate interfaces and tubulin chains comprising interface 1 and tau as interface 2. The resulting structures were compared to the lowest scoring structure to determine the built model RMSD using the Rosetta scoring function. All model building and analysis were performed on UTSW’s BioHPC computing cluster. All plots were generated with GraphPad Prism. Images were created using Pymol.

### Statistics and Reproducibility

All statistics were calculated using GraphPad Prism v8.0-10.0. Three independent ThT experiments were run for each condition. The data were normalized to the highest amplitude, and averages and standard deviations were plotted. Plots were fitted to a non-linear regression model, from which t_1/2_ values were derived. t_1/2_ error represents a 95% CI. TEM grids of endpoint ThT samples were screeded and imaged at least 4 times to obtain representative images. Flow cytometry cell aggregation was conducted in three independent experiments, whose values are plotted. Error bars represent 95% CI.

## Data availability

All MD, NMR (peptide and tauRD), ThT, XL-MS, MT stabilization, MT: tau modeling, and cell-based aggregation data are available as Source Data 1, Source Data 2, Source Data 3 and 4, Source Data 5, Source Data 6, Source Data 7, and Source Data 8, respectively. Source Data are also available on zenodo under accession number 13821565 [https://doi.org/10.5281/zenodo.13821565]. PDB IDs used in the study include 2MZ7 [https://doi.org/10.2210/pdb2MZ7/pdb], 6CVN [https://doi.org/10.2210/pdb6CVN/pdb], and 8Q92 [https://doi.org/10.2210/pdb8q92/pdb]. Supplementary Data for raw MD trajectory files, structure files for MSM modeling and validation file, and MT:tau modeling is available on zenodo under accession number 13821565 [https://doi.org/10.5281/zenodo.13821565]. All mass spectrometry data has been deposited in MassIVE under accession number MSV000093880 and is assigned to ProteomeXchange under accession number PXD048689.

## Code availability

All MD simulations were performed and analyzed using Gromacs v5.0.4 (available at http://www.gromacs.org). Markov State Model analysis was performed using pyEMMA v2.5.7 (http://emma-project.org). NMR spectra were processed, assigned, and visualized using NMRpipe (https://www.ibbr.umd.edu/nmrpipe/install.html), TopSpin (https://www.bruker.com/en/products-and-solutions/mr/nmr-software/topspin.html), Cara v1.9.1 (http://cara.nmr.ch/doku.php), ccpNMR (https://ccpn.ac.uk/software/), and Sparky 3 (https://www.cgl.ucsf.edu/home/sparky/). XL-MS data was analyzed using xQuest v2.1.5 (https://gitlab.ethz.ch/leitner_lab/xquest_xprophet). All MT:tau simulations were performed using Rosetta v3.12 (available at https://www.rosettacommons.org).

## Acknowledgments

L.A.J. is supported by an Effie Marie Cain Scholarship in Medical Research and by grants from the NIH (R01AG076459), the Chan Zuckerberg Initiative (CZI) Collaborative Science Award (2018-191983), and the Welch Foundation (I-1928-20200401 to L.A.J and I-1304 to J.R). N.E.vN. was supported by TargetALS NI-2023-NAI-S6 S.B. is supported by an NIH F31 grant from the NINDS (F31NS12751301). NMR data was acquired at the Biomolecular NMR Facility at UTSW. Heteronuclear NOE experiments were performed in the “Manuel Rico” NMR Laboratory (LMR) of the Spanish National Research Council (CSIC), a node of the Spanish Large-Scale National Facility (ICTS R-LRB). Transmission electron microscopy was performed at the Electron Microscopy Core Facility at UTSW, supported by the NIH (1S10OD021685-01A1 and 1S10OD020103-01). Computational resources were provided by the BioHPC cluster supported by the Lyda Hill Department of Bioinformatics at UTSW. We would like to thank Andrew Lemoff from the UTSW Proteomics Core for his valuable insights on troubleshooting procedures in crosslinking mass spectrometry. We would like to thank Qiong Wu from the UTSW BioNMR facility for her help in the optimization and collection of NMR data. We would also like to thank Javier Oroz from the IQF-CSIC for his guidance in the hetNOE relaxation experiments. We thank all members of the Joachimiak lab, particularly Dailu Chen, Pawel M. Wydorski, and Valerie A. Perez, and members of the UTSW CAND, specifically Jaime Vaquer-Alicea, for their discussions and input on the manuscript.

## Author Contributions

S.B. and L.A.J. initiated the project. S.B. and A.W. performed MD calculations. S.B. performed NMR experiments with guidance from J.R. S.B., and N.E.V. performed NMR relaxation experiments. S.B. and R.S. performed MSM analysis. S.B. purified all proteins involved in the study. S.B. performed all peptide and protein aggregation experiments. S.B. performed TEM of tau fibrils. S.B. and D. C. performed full-length fibril formation. S.B. and P.M.W. performed the XL-MS analysis. S.B., V.A.P. and A.W. carried out the cell-based aggregation experiments. S.B. performed the Rosetta MT:tau structure calculations. Finally, S.B. and L.A.J. conceived of and directed the research and wrote the manuscript. All authors contributed to the revisions of the manuscript.

