## supplemental information for "Amyloid-motif-dependent tau self-assembly is modulated by isoform sequence context"

**Source Data 1. Raw ThT data.**

**Source Data 2. Raw NMR data assignments (peptides)**

**Source Data 3. Raw MD and MSM trajectory analysis data**

**Source Data 4. Raw XL-MS data.**

**Source Data 5. Raw NMR data, assignments, and hetNOE intensities (tauRD)**

**Source Data 6. tauRD HSQC assignment visualization and mutation strip plots (searchable)**

**Source Data 7. Raw MT stabilization data and MT:tau modeling data.**

**Source Data 8. Raw cell-based aggregation data.**

#### SUPPLEMENTARY TABLES

| | WT R1R3 | | | P270S R1R3 | | |
|---|---|---|---|---|---|---|
| Residue | Random Coil (RC) | solution NMR 10°C | solution NMR 25°C | Random Coil (RC) | solution NMR 10°C | solution NMR 25°C |
| THR_HA | 4.25 | 4.23 | 4.26 | 4.25 | 4.22 | 4.26 |
| GLU_HA | 4.18 | 4.22 | 4.25 | 4.18 | 4.22 | 4.24 |
| ASN_HA | 4.56 | 4.63 | 4.66 | 4.56 | 4.63 | 4.65 |
| LEU_HA | 4.25 | 4.25 | 4.28 | 4.25 | 4.25 | 4.27 |
| LYS_HA | 4.14 | 4.19 | 4.20 | 4.14 | 4.18 | 4.21 |
| HIS_HA | 4.52 | 4.59 | 4.60 | 4.52 | 4.55 | 4.59 |
| GLN_HA | 4.51 | 4.59 | 4.61 | 4.26 | 4.33 | 4.36 |
| PRO/SER_HA | 4.33 | 4.39 | 4.40 | 4.38 | 4.42 | 4.45 |
| GLY_HA1 | 3.95 | 3.94 | 3.95 | 3.99 | 3.99 | 3.93 |
| GLY_HA2 | 3.95 | 3.98 | 3.92 | 3.99 | 3.96 | 3.92 |
| GLY_HA1 | 3.98 | 3.90 | 3.93 | 3.98 | 3.94 | 3.94 |
| GLY_HA2 | 3.98 | 3.95 | 3.91 | 3.98 | 3.90 | 3.92 |
| GLY_HA1 | 3.97 | 3.91 | 3.92 | 3.97 | 3.93 | 3.93 |
| GLY_HA2 | 3.97 | 3.95 | 3.92 | 3.97 | 3.90 | 3.93 |
| LYS_HA | 4.26 | 4.31 | 4.34 | 4.26 | 4.30 | 4.34 |
| VAL_HA | 3.94 | 4.02 | 4.06 | 3.94 | 4.03 | 4.07 |
| GLN_HA | 4.25 | 4.32 | 4.35 | 4.25 | 4.31 | 4.35 |
| ILE_HA | 4.09 | 4.03 | 4.08 | 4.09 | 4.03 | 4.08 |
| VAL_HA | 3.98 | 4.02 | 4.05 | 3.98 | 4.02 | 4.05 |
| TYR_HA | 4.49 | 4.51 | 4.55 | 4.49 | 4.50 | 4.54 |
| LYS_HA | 4.48 | 4.17 | 4.18 | 4.48 | 4.16 | 4.20 |
| | WT R2R3 | | | P301S R2R3 | | |
| Residue | Random Coil (RC) | solution NMR 10°C | solution NMR 25°C | Random Coil (RC) | solution NMR 10°C | solution NMR 25°C |
| LYS_HA | 4.23 | 4.19 | 4.23 | 4.23 | 4.19 | 4.23 |
| GLU_HA | 4.47 | 4.56 | 4.58 | 4.47 | 4.55 | 4.57 |
| ASN_HA | 4.61 | 4.64 | 4.66 | 4.61 | 4.64 | 4.65 |
| ILE_HA | 4.06 | 4.06 | 4.09 | 4.06 | 4.05 | 4.08 |
| LYS_HA | 4.19 | 4.25 | 4.29 | 4.19 | 4.25 | 4.27 |
| HIS_HA | 4.58 | 4.59 | 4.62 | 4.58 | 4.59 | 4.61 |
| VAL_HA | 4.27 | 4.35 | 4.38 | 4.02 | 4.10 | 4.12 |
| PRO/SER_HA | 4.33 | 4.34 | 4.36 | 4.39 | 4.42 | 4.45 |
| GLY_HA1 | 3.95 | 4.01 | 4.02 | 3.99 | 4.01 | 4.02 |
| GLY_HA2 | 3.95 | 3.91 | 3.94 | 3.99 | 3.99 | 3.98 |
| GLY_HA1 | 3.98 | 3.96 | 3.98 | 3.98 | 3.98 | 4.02 |
| GLY_HA2 | 3.98 | 3.97 | 3.97 | 3.98 | 3.97 | 3.98 |
| GLY_HA1 | 4.01 | 3.95 | 3.98 | 4.01 | 3.97 | 3.97 |
| GLY_HA2 | 4.01 | 3.94 | 3.96 | 4.01 | 3.96 | 3.96 |
| SER_HA | 4.41 | 4.47 | 4.49 | 4.41 | 4.46 | 4.48 |
| VAL_HA | 3.98 | 4.09 | 4.12 | 3.98 | 4.09 | 4.12 |
| GLN_HA | 4.25 | 4.3 | 4.33 | 4.25 | 4.3 | 4.33 |
| ILE_HA | 4.09 | 4.02 | 4.07 | 4.09 | 4.02 | 4.07 |
| VAL_HA | 3.98 | 4.02 | 4.05 | 3.98 | 4.02 | 4.05 |
| TYR_HA | 4.49 | 4.5 | 4.55 | 4.49 | 4.5 | 4.53 |
| LYS_HA | 4.48 | 4.16 | 4.2 | 4.48 | 4.17 | 4.2 |

**Supplementary Table 1. Cα proton chemical shifts (Hα) for random coil, experimental NMR at 10°C and 25°C.**

#### SUPPLEMENTARY FIGURES and LEGENDS

Supplementary Figure 1

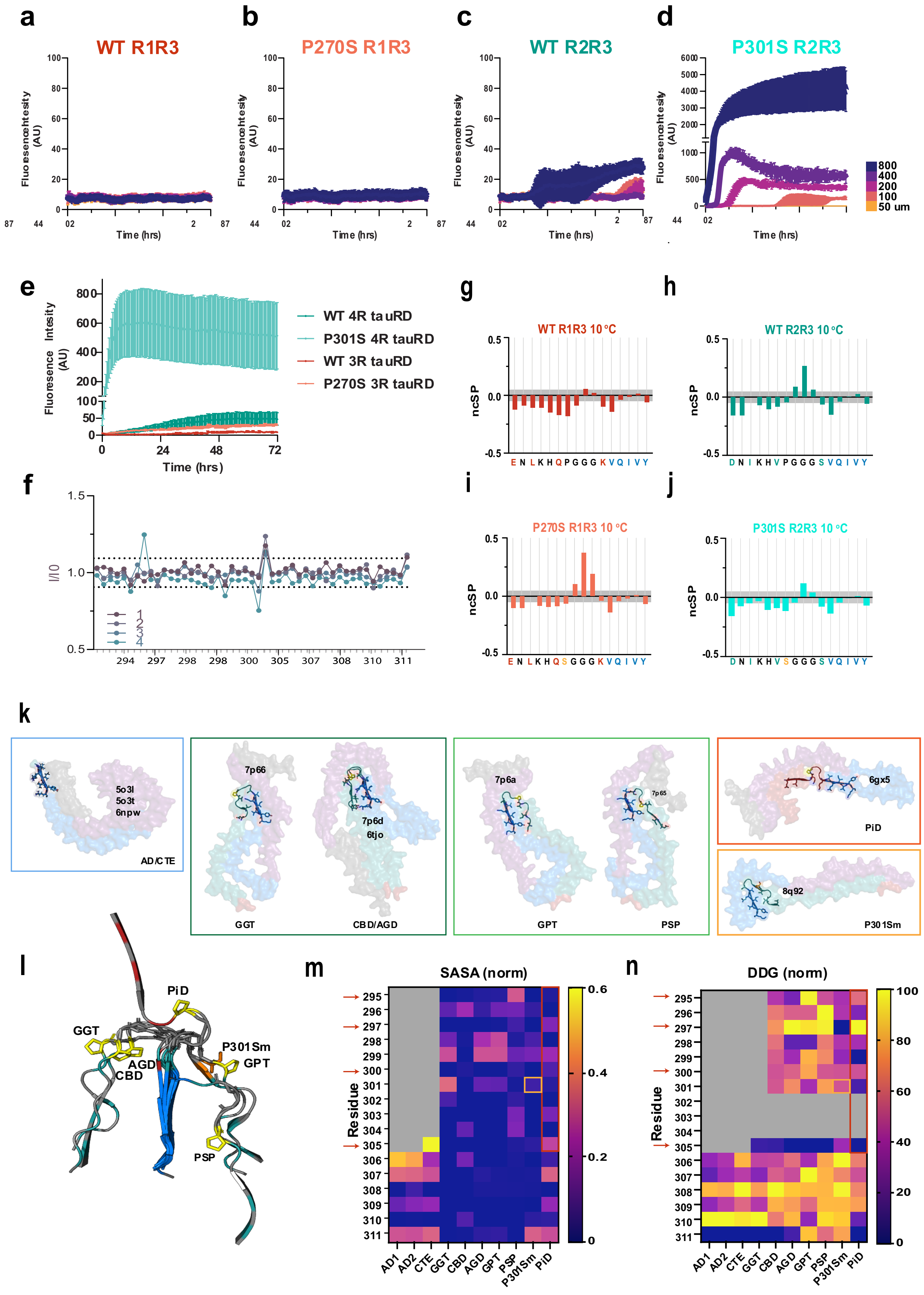

**Supplementary Figure 1. Changes in the conformation of tau fragments correlate with aggregation propensity. a-d**. ThT fluorescence aggregation assay of peptides WT $R1R3_{263\text{-}311\Delta R2}$ (**a**), P270S $R1R3_{263\text{-}311\Delta R2}$ (**b**), WT $R2R3_{295\text{-}311}$ (**c**), and P301S $R2R3_{295\text{-}311}$ (**d**) at 800, 400, 200, 100, and 50 µM (navy to orange) incubated at 37°C with intermittent shaking for 72 hrs. The data are presented as an average + /− SD for n = 3 replicates. **e.** ThT fluorescence aggregation assay of 4R WT tauRD (green), P301S 4R tauRD (cyan), 3R WT tauRD (red), and P270S 3R tauRD (orange) at 10 µM with 2X heparin concentration incubated at 37°C with intermittent shaking for 72 hrs. The data are presented as an average + /− SD for n = 3 replicates. **f.** P301S $R2R3_{295\text{-}311}$TOCSY spectra were measured four additional times for 10 hrs each, with peak intensities normalized to the first measurement across all assigned peaks for the five spectra. Data is colored from purple to blue for each timepoint. **g-j.** Neighbor corrected structure propensity (ncSP) calculated from peptide 10˚C Hα chemical shift assignments using the ncSPC calculator colored as in Figure 1d-g. **k.** Surface representation of the tau fibril assemblies from AD, CTE, GGT, CBD, AGD, GPT, PSP, PiD and P301Sm. Structures are colored by repeat domain, as in Fig. 1a. The region 295-311 (263-311 for PiD) is shown in cartoon representation with the unique R1/R2 residues and amyloid motif residues shown as sticks. Structures are grouped by the orientation of the 295-311 (263-311 for PiD) interactions: disordered (blue), "-Q-V-K" face (dark green), "V-I-Y" face (light green), extended – 3R-derived (red), mutant P301S derived (orange). Structures are labeled according to tauopathy (acronym described in methods) **l.** Overlay of 295-311 (263-311 for PiD) elements across selected structures, colored as in Fig. 1a, P301/S301 shown in sticks (yellow/orange). **m.** Solvent accessible surface area (SASA), normalized to maximum SASA per residue (purple to yellow), for 295-311 region and equivalent region in PiD (263-311). **n.** In silico rosetta $\Delta REU_{mut\text{-}wt}$ calculation of stability measurements for 295-311 from Mullapudi et. al, 2023 alanine scan. $\Delta\Delta G$ values normalized to the 295-311 region represent the loss of structural stability by incorporation of alanine at each position (purple to yellow).

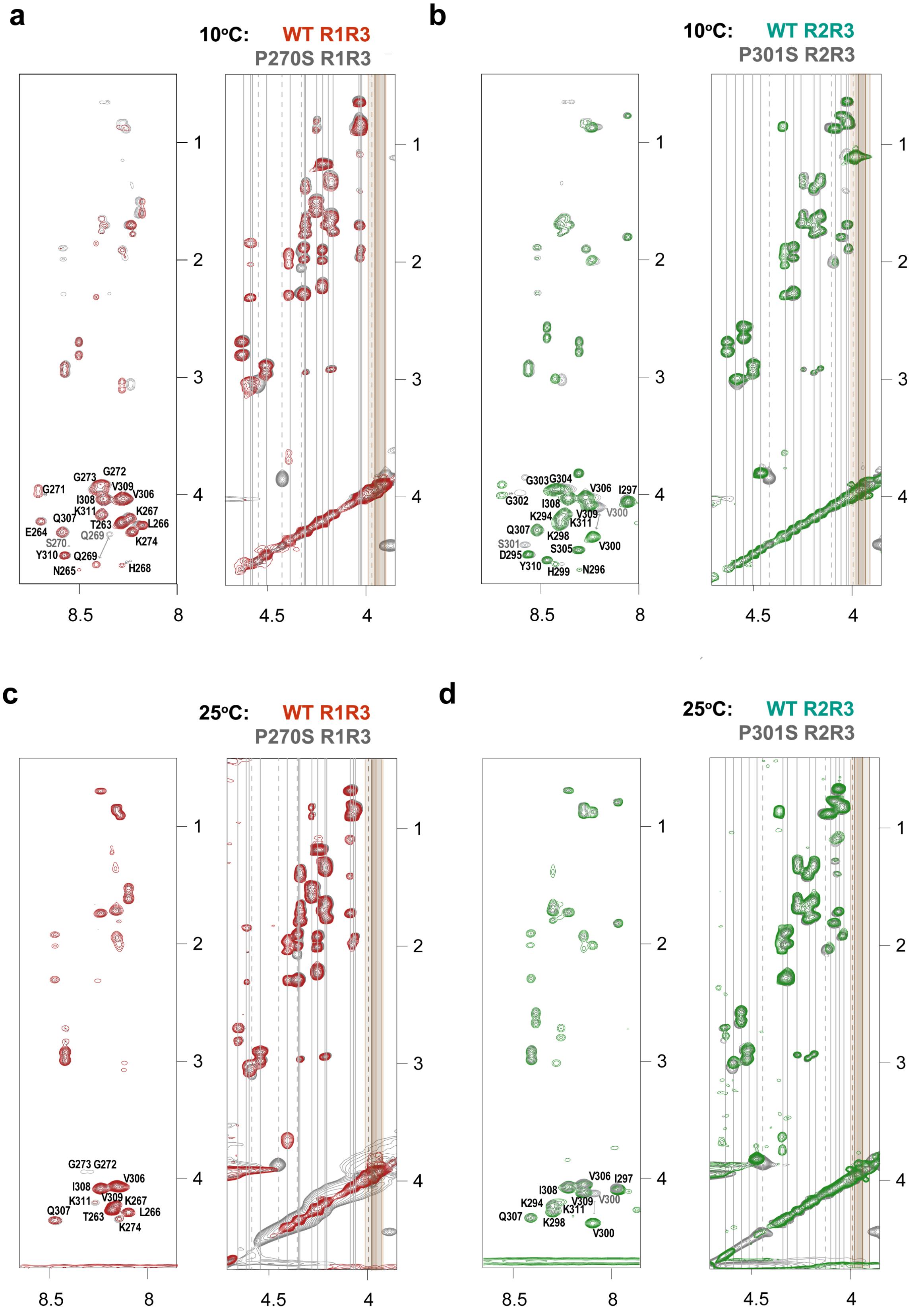


**Supplementary Figure 2. Homonuclear NMR spectral assignments. a-d.** TOCSY HN-HA and HA-HA+SC region spectra from peptide homonuclear NMR of WT R1R3$_{263\text{-}311\Delta R2}$ (red) and P270S R1R3$_{263\text{-}311\Delta R2}$ (gray) at 10˚C (**a**) and 25˚C (**c**), and of WT R2R3$_{295\text{-}311}$(green) and P301S R2R3$_{295\text{-}311}$ (gray) at 10˚C (**b**) and 25˚C (**d**). NH-HA peak assignments are labeled, proline to serine mutant peaks are labeled in gray, HA assignments are marked by vertical lines with glycine HA1 and HA2 labeled by brown bars, and proline to serine mutant shifts are marked by dashed lines.

#### Supplementary Figure 3

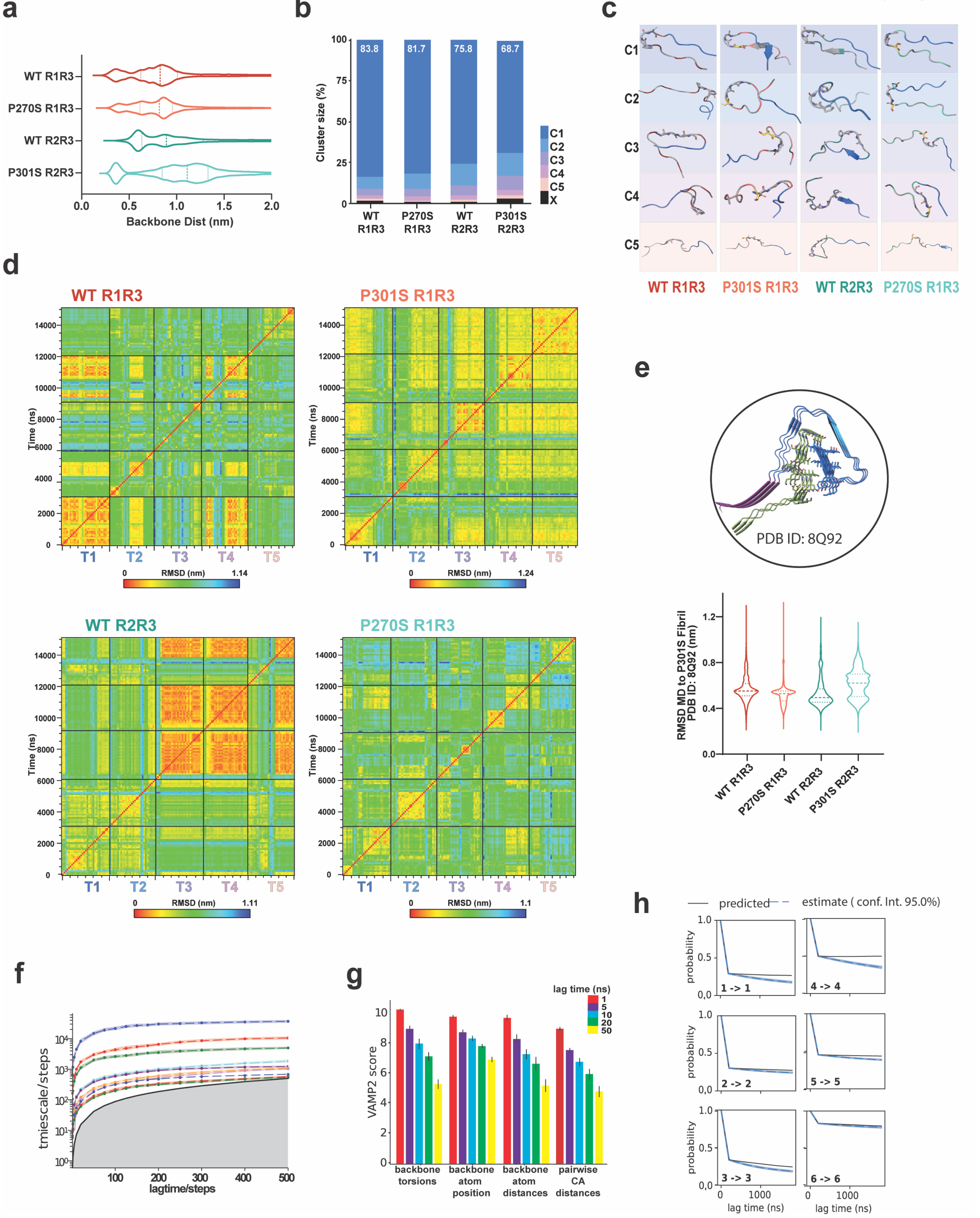

**Supplementary Figure 3. Vertification of sampling and validation of MD simulations with MSM models a.** Comparison of first step MD cumulative ensembles for WT R1R3$_{263\text{-}311\Delta R2}$ (red), P270S R1R3$_{263\text{-}311\Delta R2}$ (green), WT R2R3$_{295\text{-}311}$ (orange), and P301S R2R3$_{295\text{-}311}$ (cyan) by backbone distance between the center of mass of the N-term residues 1-6 (TENLKH/KDNIKH) and C-term amyloid motif residues (VQIVYK). The data is shown as violin plots with median and 25/75% quartiles labeled in dashed lines. **b.** Bar plots illustrating the cluster size for the production stage i MD ensembles according to a 0.54nm cutoff. The clusters for each ensemble are colored blue to pink for clusters 1-5, with the remaining clusters colored black. **c.** Initial structures for second step MD simulations were obtained from the mean structure of the top 5 clusters in step 1 (background color). Structures shown in cartoon representation with 'VQIVYK' colored in blue, unique residues are colored as in Fig. 1b, and similar residues are colored in gray. **d.** 2-dimensional root means squared deviation (RMSD) matrix between all five 3μs simulations (T1-T5) from the second step of MD for peptides WT R1R3$_{263\text{-}311\Delta R2,}$ P270S R1R3$_{263\text{-}311\Delta R2,}$ WT R2R3$_{295\text{-}311,}$ and P301S R2R3$_{295\text{-}311}$. Data is colored in a rainbow scale based on RMSD in nm. Matrix labels (black line) distinguish separate simulations**. e.** RMSD of 15 μs MD ensemble to the P301Sm tau fibril structure (PDB ID 8Q92, top). WT R1R3 (red), P270S R1R3 (orange), WT R2R3 (green), and P301S R2R3 (cyan). The data are shown as violin plots with median and quartiles labeled in dashed lines. **f.** Convergence behavior for the implied timescale to obtain MSM lag time. The timescale shown for the 10 slowest processes is colored blue to orange. The gray area dictates the timescale horizon. **g**. Comparison of VAMP2 scores for different feature selection models for lag times of 1ns (red), 5ns (purple), 10 ns (blue), 20 ns (green), and 50 ns (yellow). **h.** CK test results for combined MSM for transitions within each state are shown as MSM model maximum likelihood estimate in black and a Bayesian mean with the 95% confidence interval in blue.

**Supplementary Note 1**

The initial structure for the four peptides (WT R2R3$_{295\text{-}311}$, P301S R2R3$_{295\text{-}311}$, WT R1R3$_{263\text{-}311\Delta R2,}$ and P270S R1R3$_{263\text{-}311\Delta R2}$) were built as extended peptide chains that were minimized in GROMACS, for production stage i. Minimized systems were run in 5 independent 1 µs simulations per peptide (Fig. 2a,- stage i). The independent trajectories were concatenated, and from the resultant ensemble, we clustered all conformations using a cutoff of 0.54nm(Fig. 2a, stage ii). The largest cluster adopted a beta-hairpin-like conformation for all peptides (Supplementary Fig. 3b, C1 structures), yet P301S R2R3$_{295\text{-}311}$ had the smallest top cluster, 68.7% (Supplementary Fig. 3b). To sample the stability of a wider range of states, we implemented a second production stage of longer simulations initiated from the mean structure of the top 5 clusters for the four peptides (Supplementary Fig. 3c). Minimized systems were simulated for 3 µs using TIP3 water and 150mM NaCl in the solvent (Fig. 2a, stage iii). The 2-D root mean squared deviation (RMSD) was measured across 5 individual trajectories, revealing heterogeneous sampling across most trajectories independent of initial structure for most trajectories (Supplementary Fig. 3d). We note the exception of restricted sampling in T1 of WT R1R3 and in T3 and T4of WT R2R3 where we observe restricted sampling. However, we note that for WT R1R3, the state is also reached from the initial structure of T4 and, in that trajectory, can exchange to distinct states, while in WT R2R3, this state is reached from distinct initial structures and is accessible from the unfolded initial state of T5 suggesting a relevant sampling restriction.

As we aim to understand states that are on pathway to the fibril state, we compared the MD ensembles to a recently published cryo-EM structure of tau fibrils isolated from tauopathy mice encoding P301S tau (PDB: 8Q92). The structural comparison highlights that the ensembles of WT R2R3295-311 and P301S R2R3295-311 sample conformations are consistent with the turn observed in the fibril (Supplementary Fig. 3g, rmsd <0.4). Interestingly, the P301S R2R3295-311 has a bimodal RMSD distribution, suggesting a fraction of the ensemble is compatible with the fibrillar fold while the other population of structures is more expanded (Supplementary Fig. 3g).

The MSM model was built using a pooled ensemble of the trajectories of all four peptides (Fig 2a, stage iv-iv). We used the minimum pairwise distance between residues 1-8 against amyloid motif residues 12-18 for discretization. To build the MSM model we used the k-means algorithm with 200 cluster centers and selected maximum convergence time of 20 ns (200 steps) after visualizing the implied timescales (Supplementary Fig. 4a). We assessed our selected featurization parameter and lag time by comparing VAMP2 scores and observing least dependence of feature selection at 20ns lag time (Supplementary Fig. 4b). We determined the number of metastable states is 6 according to the PCCA+ algorithm. Additionally, we built MSM models for the final concatenated trajectories of each peptide (WT R1R3, P270S R1R3, WT R2R3, P301S R2R3) and assessed the validity of the 20ns lag time and featurization as for the combined model. Finally, we validated the selection of 6 metastable states using the C-K test for the combined and individual peptide models (Supplementary Fig. 4c).

#### Supplementary Figure 4

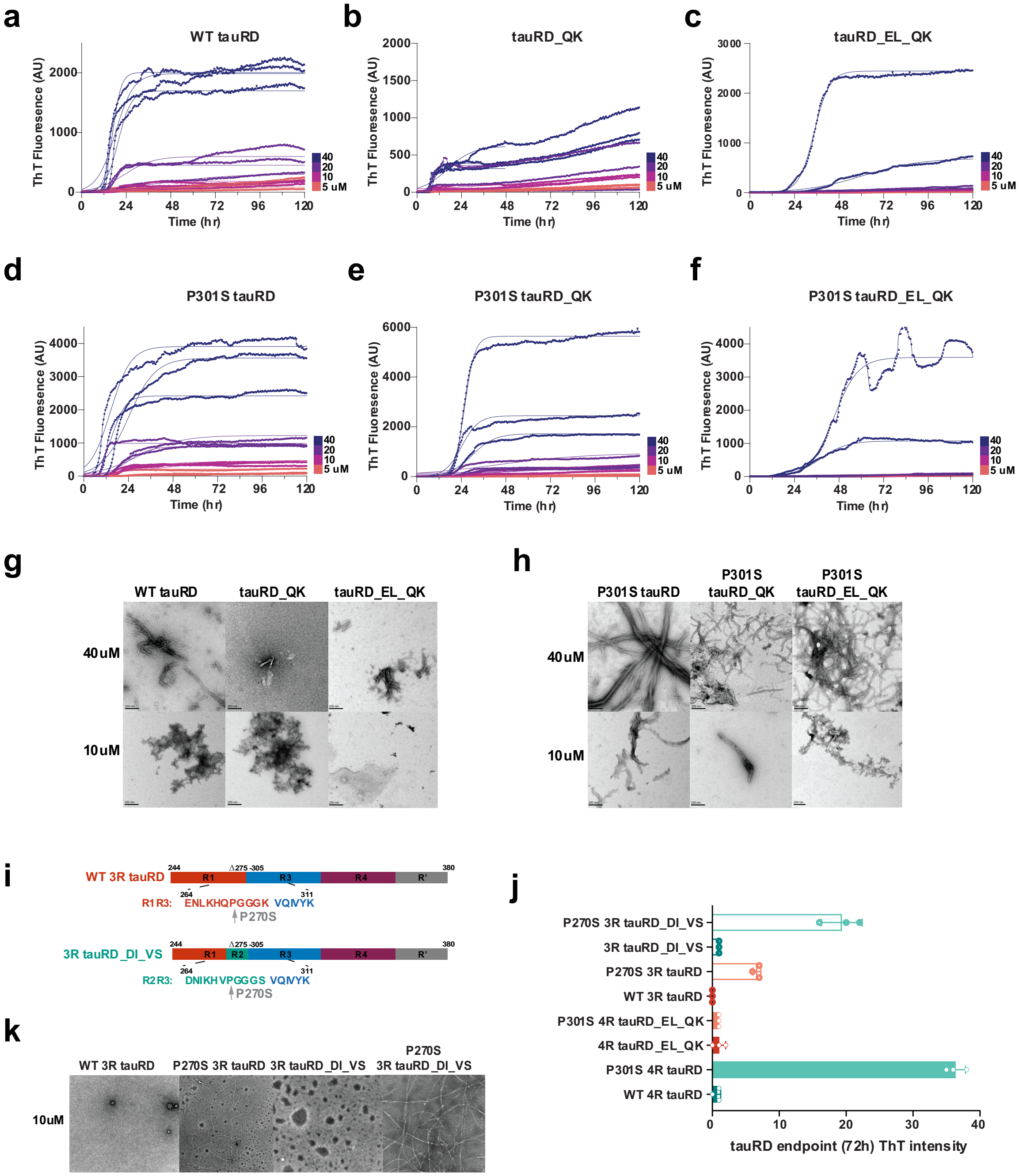


**Supplementary Figure 4. Designed tau mutations in proximity to amyloid motifs reduce tauRD aggregation in vitro. a-f.** ThT fluorescence aggregation assay for 4R tauRD WT (a), tauRD_QK (b), tauRD_EL_QK (c), P301S (d), P301S tauRD_QK (e), and P301S tauRD_EL_QK (f). Constructs at 40 μM, 20 μM, 10 μM, and 5 μM (navy to orange) mixed with 2X heparin and allowed to aggregate in the presence of ThT at 37˚ C with intermittent shaking every 30 minutes for 7 days. Constructs were tested in n=3 biological replicates, and resultant curves were fit using a non-linear regression model in GraphPad Prism. **g-h.** Representative TEM images of 120 hrs ThT endpoints for each construct at 10 μM and 40 μM. **i.** Cartoon schematic of 3R tauRD designed construct with isoform switch mutations with and without P270S mutation. **j.** ThT fluorescence intensity of aggregation endpoints (72 hrs) for 3R tauRD constructs (outline): WT (red), P270S (orange), tauRD_DI_VS (green), and P270S tauRD_DI_VS (cyan) and 4R tauRD constructs (filled) colored as in figure 4b. Samples at 10uM with 2X heparin. **k.** Representative TEM images of 3R tauRD aggregation endpoints.

### Supplementary Figure 5

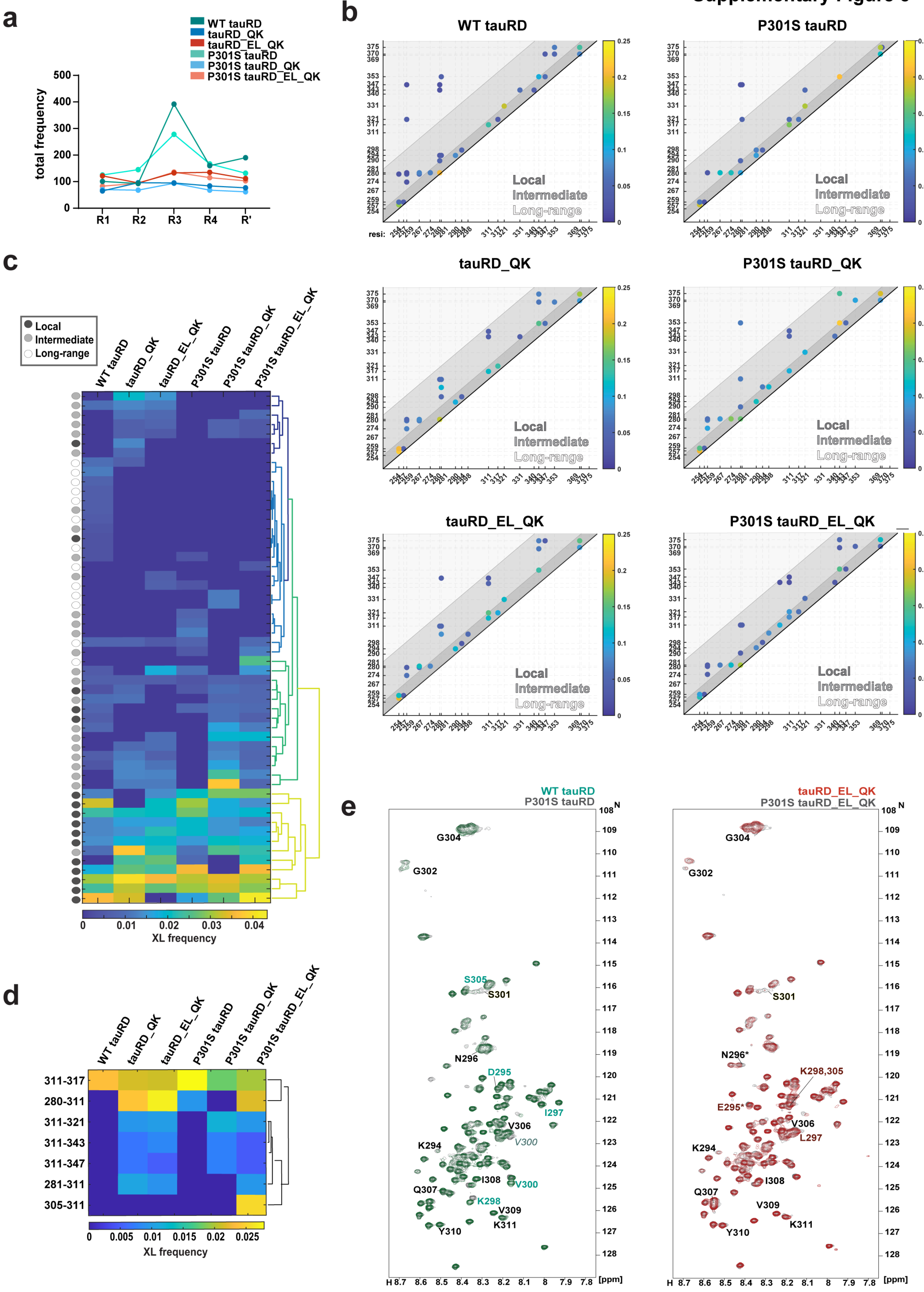

**Supplementary Figure 5. Balance of intermediate and long-range contacts correlates with reduced aggregation propensity. a.** Mono-link modification frequency of designed constructs grouped by repeat: R1 (244-274), R2 (275-305), R3 (306-336), R4 (337-368), and R' (369-380) for WT tauRD (teal), tauRD_QK (blue), tauRD_EL_QK (red), P301S tauRD (cyan), P301S tauRD_QK (light blue) and P301S tauRD_EL_QK (orange). **b.** Scatter plots of cross-link and loop-links are shown as contact maps. Pairs of cross-links are shown as dots and are colored by their normalized frequency across 5 technical replicates. The contact order is indicated by distance to help distinguish contacts within repeats (local; < 10 residues), between adjacent repeats (intermediate; 11-39 residues), and long-range interactions (> 40 residues) and is indicated by white, light grey and grey colors, respectively. **c.** Dendrogram produced by hierarchical clustering using Ward's method of cross-link pair frequencies. The color scale is colored in plasma from yellow (highest cross-link frequency) to blue (lowest cross-link frequency) for all identified cross-link pairs. Each cross-link is labeled as local (dark gray), intermediate (gray), and long-range (light gray). **d.** Dendrogram produced as in d for cross-links to position 311. **e.** HSQC spectra for WT tauRD, green, and P301S tauRD, gray (left), and tauRD_EL_QK, red, and P301S tauRD_EL_QK, gray (right). Assignments for residues in the 294-311 fragment labeled (Full Assignments in Source Data 5).

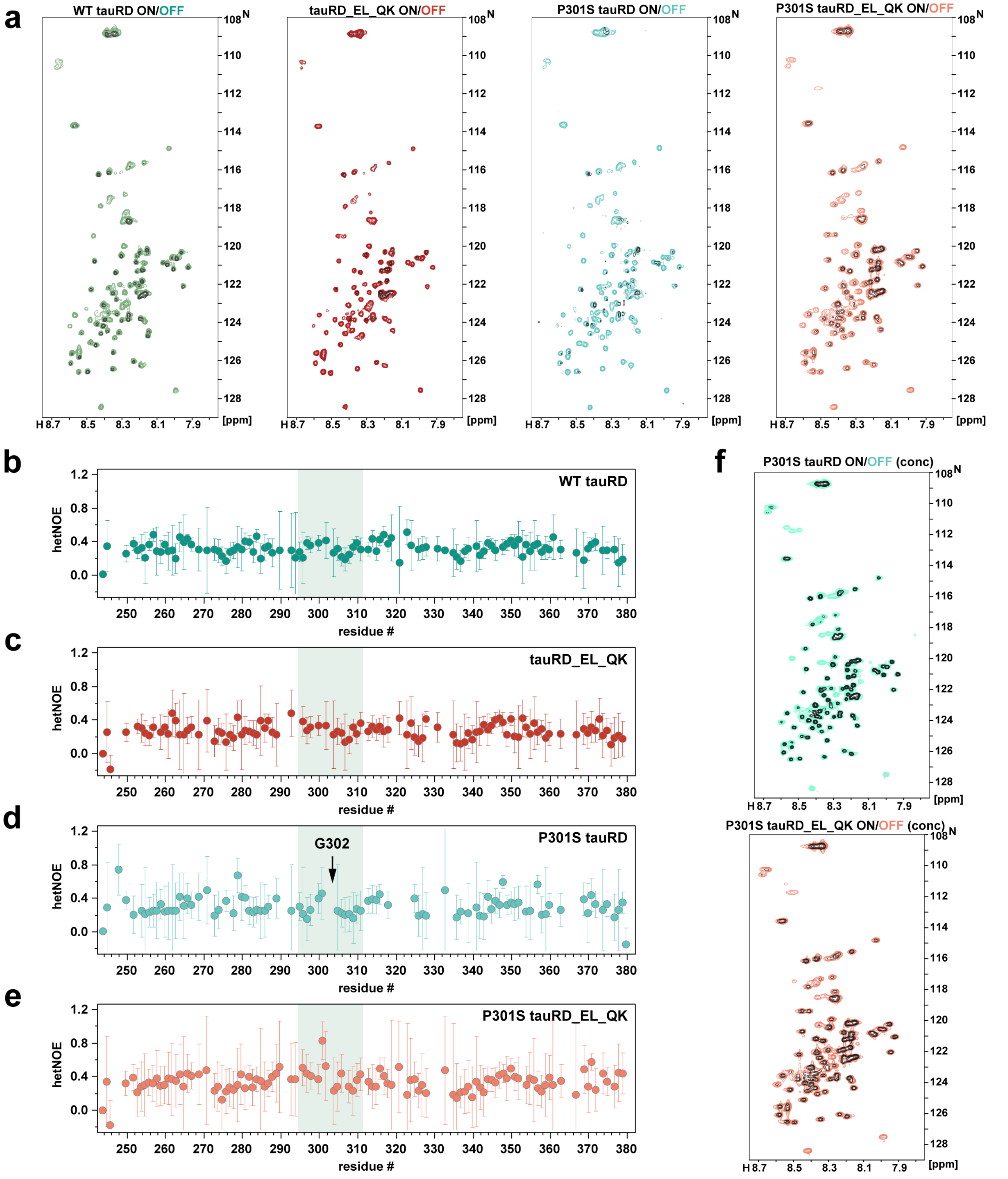


**Supplementary Figure 6. Measuring dynamics using steady-state hetNOE experiments. a.** $^{15}$N-$^{1}$H HSQC spectra tor tauRD constructs using the $^{15}$N-$^{1}$H heteronuclear NOE experiment at ~150 uM. Reference spectra are green, red, cyan, and orange for WT, tauRD_EL_QK, P301S, and P301S tauRD_EL_QK. Saturation experiment spectra for each construct overlayed in gray (black(+)/gray(-)). (**b-e**) $^{15}$N-$^{1}$H heteronuclear NOE experiments for WT, tauRD_EL_QK, P301S, and P301S tauRD_EL_QK colored in green, red, cyan, and orange, respectively. The ratio is calculated by comparing the integrated peak volume between the saturation and reference spectra. Peak errors were determined from the standard deviation of the intensity distribution in signal-less spectral regions. **f**. $^{15}N$-$^{1}H$ HSQC spectra for P301S and P301S Full mutant tauRD using the $^{15}N$-$^{1}H$ heteronuclear NOE experiment at high concentration (300μM). Reference spectra, colored cyan and orange for P301S and P301S tauRD_EL_QK, respectively, and saturation spectra, colored as in c.

#### Supplementary Figure 7

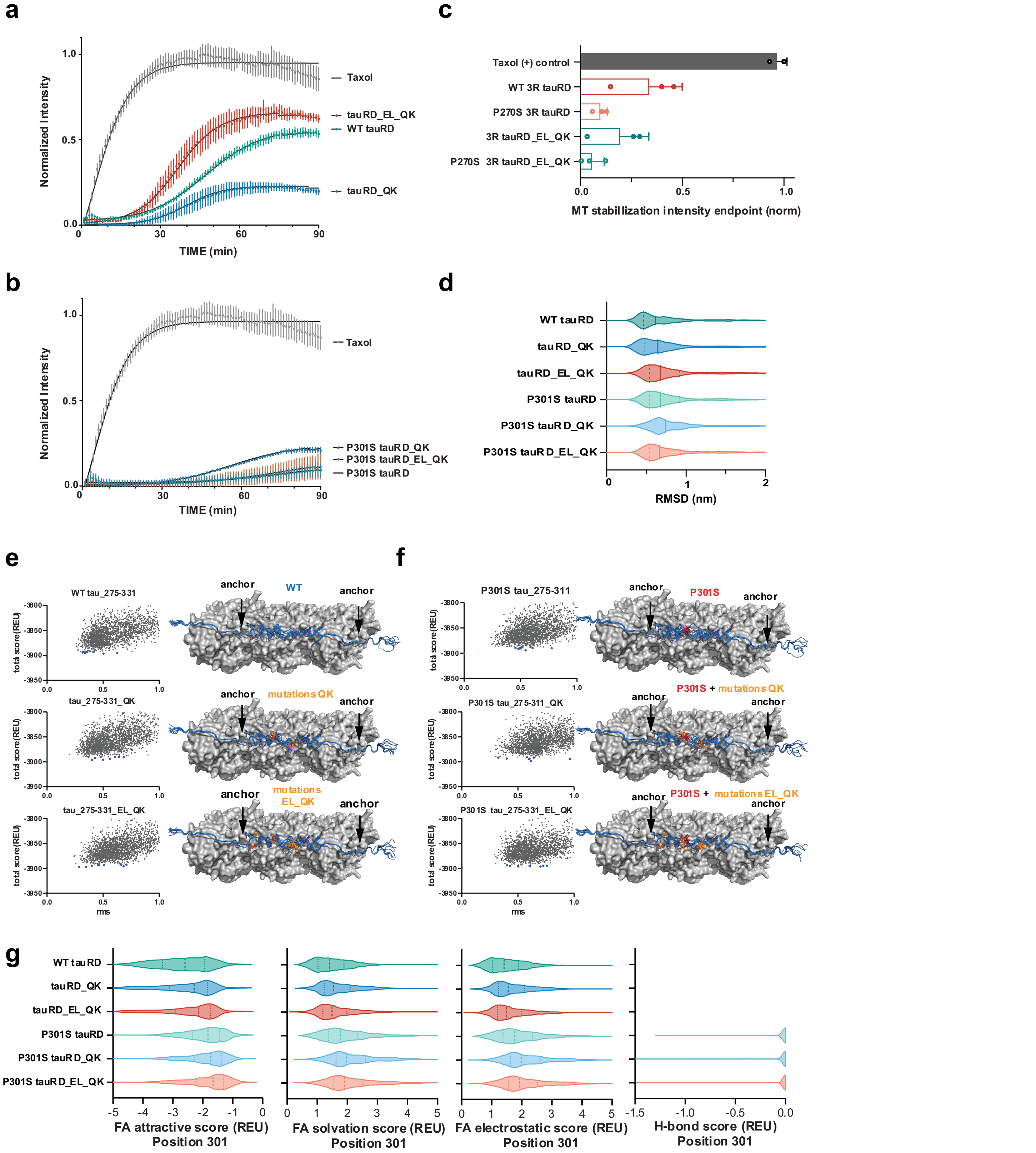


**Supplementary Figure 7. Designed tau sequences maintain MT binding but cannot overcome the effect of P301S. a-b.** MT stabilization assay comparing (a) WT (green), tauRD_QK (blue), and tauRD_EL_QK (red) and (b) P301S (cyan), P301S tauRD_QK (light blue), and P301S tauRD_EL_QK (orange). Taxol as control (gray) was the same for a-b. Data are shown as averages with standard deviation across three replicates. **c**. MT stabilization assay comparing normalized endpoint intensity of WT 3R tauRD (red), P270S 3R tauRD (orange), 3R tauRD_DI_VS (green), P270S 3R tauRD_DI_VS (cyan), and Taxol as a control (gray). **d.** RMSD distributions of tau conformations bound to MTs for WT (green), tauRD_QK (blue), tauRD_EL_QK (red), P301S (cyan), P301S tauRD_QK (light blue), and P301S tauRD_EL_QK (orange). The data are shown as violin plots with median and quartiles labeled in dashed lines. **e-f**. Scatter plots illustrating the distribution of MT:tau energies (Total Score, REU) and RMSD to the lowest scoring model. The 10 lowest-scoring models of MT:tau are shown. Tau is shown in a cartoon, and MT is shown in a surface representation. MT is colored grey. Tau is colored in blue, and D295E, I297L, V300Q, and S305K mutation sites are colored in orange. The P301S site is colored in red. MT:tau anchor points are indicated by an arrow. **g**. Rosetta energy terms for position 301: full atom attractive, full atom solvation, full atom electrostatic, and hydrogen bond score are constructs colored as in d. The data are shown as violin plots with median and quartiles labeled in dashed lines.

Supplementary Figure 8

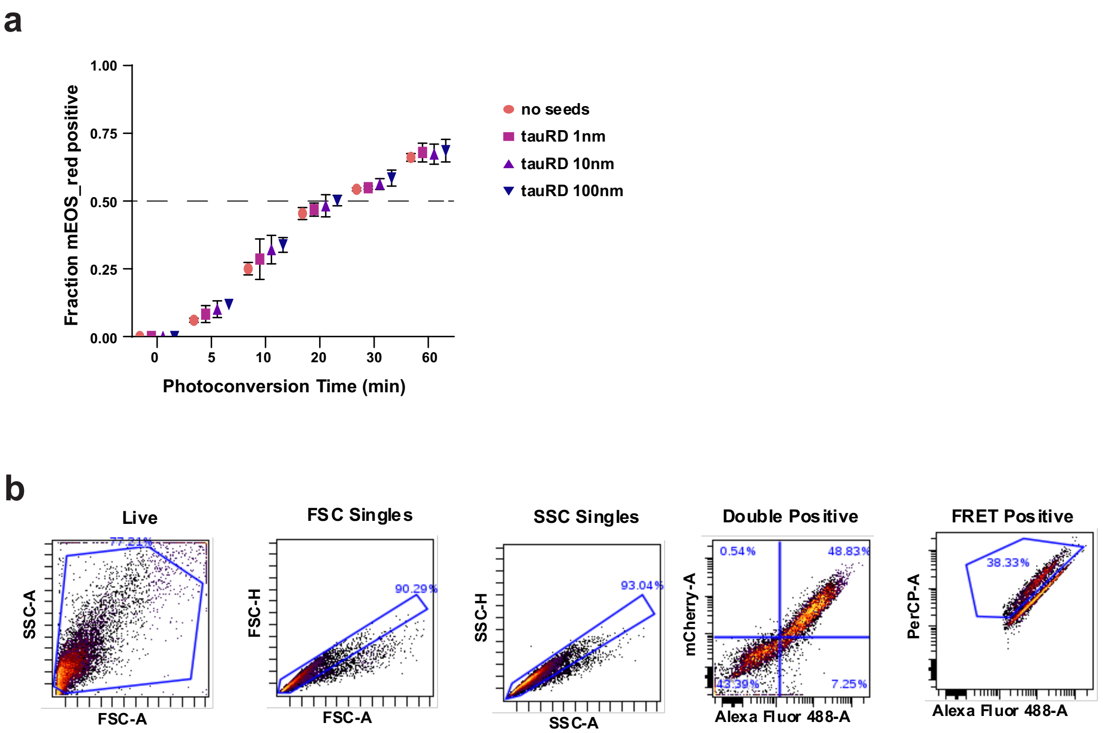


**Supplementary Figure 8. Flow cytometry gating strategy is used to quantify the percentage of cells with FRET positive signal. a.** Quantification of mEOS3.2 photoconversion as a function of time and colored by the concentration of WT tauRD fibrils used in the seeding experiment. 0, 1, 10, and 100nM tauRD fibril conditions are colored orange to blue. **b.** Gating strategy to extract live, single, Alexa-488 (green), and mCherry (red) channel double-positive cells and expression level for FRET quantification.
